# Chromatin is dispensable for bacterial life

**DOI:** 10.64898/2026.08.21.744486

**Authors:** Paul Villain, Antoine Hocher, Jacques Serizay, Chad Whilding, Alex Montoya, Pavel V Shliaha, Romain Koszul, Tobias Warnecke

## Abstract

Inside cells, DNA is intimately associated with proteins, forming chromatin. The protein constituents of chromatin vary across the tree of life: histones are the principal building blocks of chromatin in eukaryotes and many archaea, whereas bacteria typically encode a collection of nucleoid-associated proteins (NAPs) that wrap, bend, bridge or coat the DNA. Although chromatin proteins appear to be a universal feature of cellular life, DNA-templated processes such as transcription, replication, and DNA repair can take place *in vitro* in the absence of chromatin, raising the possibility that cellular systems might exist – or could be built – that lack chromatin proteins. To explore this possibility, the molecular consequences and potential systemic adjustments required for life without chromatin, we serially deleted the nine most abundant NAPs from *E. coli* (*hupA, hupB, ihfA, ihfB, hns, stpA, fis, dps, lrp*), resulting in a strain (ΔNAP9) that lacks its native chromatin. Using an array of different techniques, we document change – and sometimes surprising lack thereof – in compaction, composition and 3D architecture of the nucleoid, supercoiling, prophage activity, growth, viability, and genetic make-up of ΔNAP9. Most notably, we find that ΔNAP9 exhibits global dysregulation of gene expression, marked by a striking homogenization of transcriptional output across the genome that is reminiscent of the effects of histone depletion in eukaryotic cells. Our results reinforce the notion that chromatin plays a key role in compartmentalizing the use of genomic information, enabling both the localized suppression of selfish elements and dynamic reprogramming of genome activity in response to environmental change. At the same time, the successful construction of ΔNAP9 demonstrates that bacterial cells can carry out basic cellular functions in the absence of co-evolved chromatin proteins, highlighting the potential for radical (re-)engineering of prokaryotic chromatin and systems of gene expression.

## INTRODUCTION

In all cells we know, DNA exists in complex with abundant, basic proteins. These proteins wrap, coat, stiffen or bend the DNA, and thereby affect how genomic information can be read by the enzymes that carry out replication, transcription, DNA repair, and recombination (Luijsterburg et al. 2008). Histones are the principal example of such proteins in eukaryotes, abundant enough to parcel the majority of the genome into nucleosomes. Their presence limits access to the genome on a global scale, acting both as a throttle on and adaptive funnel for transcriptional activity (Struhl 1999; Kornberg and Lorch 2020; Madhani 2013). In prokaryotes too DNA is intimately associated with abundant DNA-binding proteins; some are phylogenetically widespread, like HU, others restricted to specific lineages (Hołówka and Zakrzewska-Czerwińska 2020). These nucleoid-associated proteins (NAPs), typically present in thousands or tens of thousands of copies per cell (Mori et al. 2021; Azam et al. 1999), collectively affect DNA-templated processes and structure the bacterial nucleoid.

Informed by the biology of both eukaryotes and prokaryotes, chromatin has come to occupy a central place in our understanding of genome regulation; so central, in fact, that it might be difficult to envisage a cellular system of gene expression that operates without dedicated chromatin proteins. Could we build such a system from scratch or remove chromatin proteins from an existing system without fatally compromising it?

In eukaryotes, dispensing with histones appears to be challenging at best, on both an experimental and evolutionary time scale. Transient depletion of histones can be induced artificially or occur naturally, for example during early development, when it is accompanied by large-scale changes in genome activity, including widespread transcription from previously cryptic promoters (Kok et al. 2025; Cheung et al. 2008; Doris et al. 2018; Kaplan 2003; Wyrick et al. 1999; Gossett and Lieb 2012; Chari et al. 2019). Attempts to permanently eliminate histones from the cell, however, by deleting the genes that encode them, have invariably resulted in cell death (Gossett and Lieb 2012; Wyrick et al. 1999; Han et al. 1988; Kim et al. 1988; Han and Grunstein 1988). In addition, all eukaryotic genomes studied to date encode a minimal set of four core histones (H2A, H2B, H3, H4) (Soo and Warnecke 2021), supporting the notion that histones are essential to eukaryotic life. The idea of building a eukaryotic cell without histones, then, appears fanciful. Not impossible, perhaps, but dependent on overcoming myriad evolved dependencies along an engineering roadmap that is currently hard to fathom.

While the entrenchment of histones in eukaryotic genome biology appears deep, what about prokaryotes? *E. coli* – the pre-eminent model for bacterial nucleoid biology – encodes a suite of NAPs that have been characterized individually in considerable detail. This includes two paralogous HU proteins, HupA and HupB, which have been implicated in multiple cellular processes including replication initiation (Dixon and Kornberg 1984), DNA repair (Kamashev and Rouviere-Yaniv 2000), regulation of supercoiling (Broyles and Pettijohn 1986), transposition of bacteriophages (Kano et al. 1989), the three-dimensional structure of the genome (Lioy et al. 2018) and transcription (Aki et al. 1996; Dillon and Dorman 2010). It also includes H-NS and its paralog StpA, which target AT-rich sequences and are thought to have evolved as a defence system against horizontally transferred, and frequently selfish, DNA (Singh and Grainger 2013; Singh et al. 2014). Removal of H-NS causes widespread changes in the 3D genome (Gavrilov et al. 2025) and leads to activation of previously silenced regions, including cryptic prophages. This is reminiscent of histone depletion in eukaryotes but with activation patterns that - reflecting the binding patterns of H-NS/StpA – are localized rather than global (Singh and Grainger 2013; Singh et al. 2014; Amemiya et al. 2021a). Several other DNA-binding proteins, notably Fis, Dps, and IHF (encoded by paralogs *ihfA* and *ihfB*) are also abundant, affect nucleoid organization and function in multifarious ways, and are commonly categorized as NAPs (Dillon and Dorman 2010).

In sharp contrast to eukaryotes, where core histone deletions result in cell cycle arrest and – ultimately – death, all the *E. coli* NAPs highlighted above can be deleted individually. This dispensability of individual NAPs has been attributed, in part, to functional redundancy, where deletion of one NAP is buffered by the continued presence (and perhaps upregulation) of another (Schwab and Dame 2025). Indeed, several *E. coli* NAPs are paralogs of each other (e.g. H-NS and StpA) or share a common fold (HU and IHF) and can substitute for each other in specific instances. For example, HU can, by virtue of a similar DNA-bending capacity, functionally replace IHF to promote opening of the origin of replication, oriC (Hwang and Kornberg 1992), or in the context of lambda recombination (Segall et al. 1994).

However, earlier work suggested that such redundancy has its limits: while some double deletions involving paralogous genes (e.g. Δ*hupA*Δ*hupB* or Δ*hns*Δ*stpA)* (Khan and Kuzminov 2017; Huisman et al. 1989; Sonden and Uhlin 1996) or even triple deletions (Δ*hupA*Δ*hupB*Δ*ihfA*) (Goshima et al. 1990; Yasuzawa et al. 1992) are viable, attempts to generate higher-order NAP deletions have failed, leading, for example, to the suggestion that H-NS becomes conditionally essential in the absence of HU and IHF (Yasuzawa et al. 1992). Retention of HU in the smallest genome of a free-living bacterium engineered to date, *Mycoplasma mycoides* JCV-syn1.0 (Hutchison et al. 2016), further seems to suggest that, while partial redundancy exists amongst NAPs, a minimal complement of chromatin proteins is essential for bacterial life.

Here, we challenge this view, reporting on the construction of an *E. coli* strain lacking its nine most abundant NAPs: *hupA, hupB, ihfA, ihfB, hns, stpA, fis, dps,* and *lrp*.

## RESULTS

### Serial deletion of nine abundant NAP genes in E. coli

NAPs are generally considered to be abundant, basic proteins that bind double-stranded DNA with limited sequence specificity (Dillon and Dorman 2010). As DNA-binding proteins exist along a continuum of abundance and specificity, however, there is no objective dividing line that separates NAPs on one side from transcription factors on the other (Dorman et al. 2020). In deciding on a catalogue of genes to delete, we were guided by prior work on NAPs in *E. coli* (Azam and Ishihama 1999; Azam et al. 1999), by what is recognized as a NAP by the community (Dillon and Dorman 2010; Hołówka and Zakrzewska-Czerwińska 2020; Amemiya et al. 2021b), and by quantitative proteomics data from exponential and stationary phase *E. coli* cultures (Figure 1C, Figure S1), which capture the abundance of NAPs and their concomitant potential to shape the structure and accessibility of the *E. coli* chromosome on a global scale (Mori et al. 2021). We settled on a target list of nine genes: *hupA, hupB, ihfA, ihfB, hns, stpA, fis, dps,* and *lrp* (Figure 1A). We did not attempt to delete genes for DNA-binding proteins that are abundant but processive (e.g. RNA polymerase subunits), exhibit enzymatic activity (e.g. topoisomerases), have higher affinity for single-stranded DNA (e.g. *ssb1*, *cspE*) or RNA (e.g. *hfq*) than for double-stranded DNA, or where high abundance is linked to specific environmental condition (e.g.*dan*) (Teramoto et al. 2010; Lim et al. 2012). Some abundant regulators of transcription or replication, like CRP or SeqA (Dorman et al. 2020), could have reasonably been included as targets but did not make the final shortlist.

**Figure 1.**
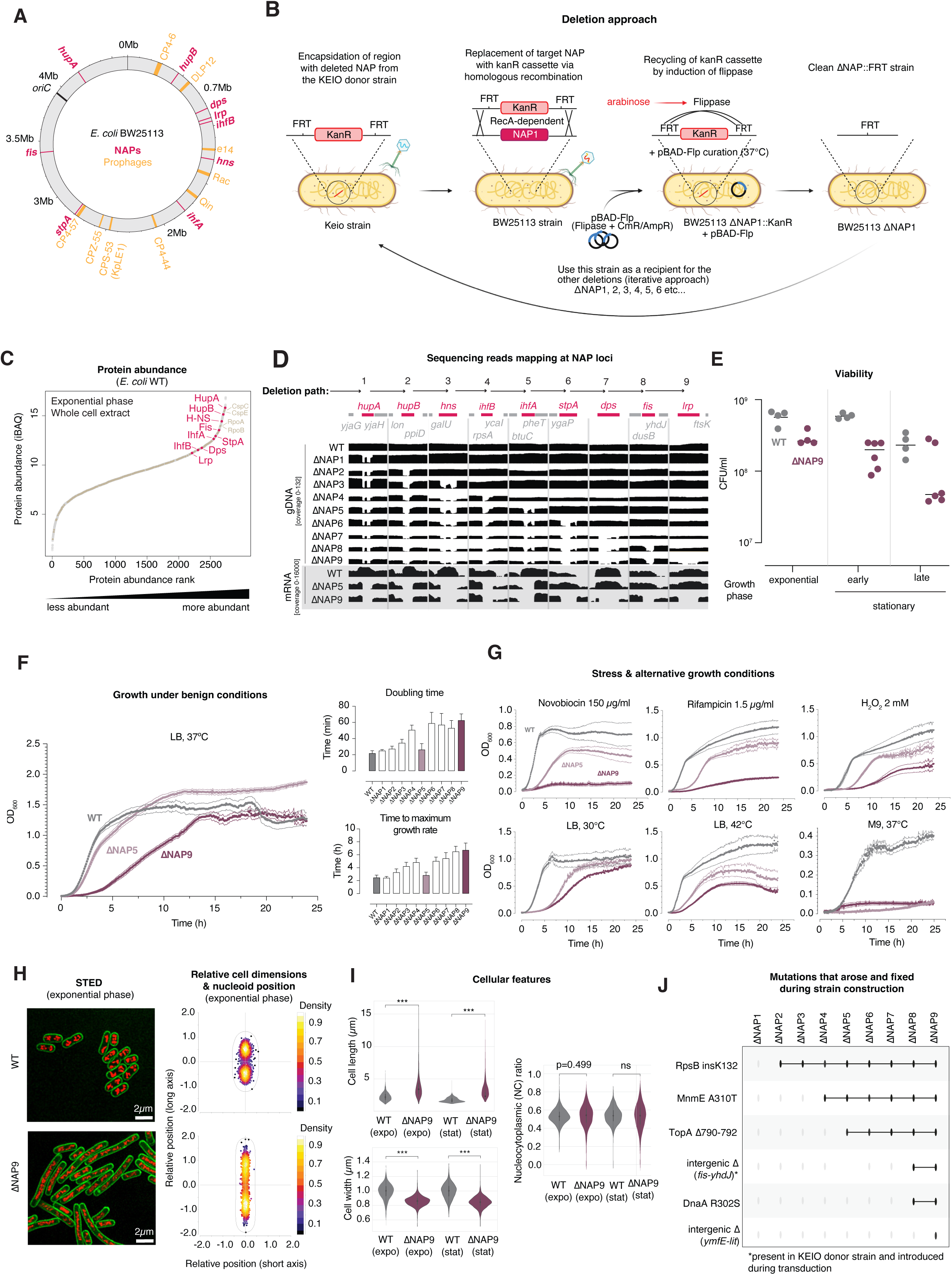
Construction and phenotyping of ΔNAP9. **A.** Location of NAPs targeted for deletion (pink) as well as cryptic prophages (orange) on the main chromosome of *E. coli* BW25113. **B.** Schematic of the deletion strategy used for the construction of ΔNAP9. Individual deletions were phage P1-transduced from the Keio collection. The KanR cassette was recycled between each round of deletion by expressing flippase from the pBAD-Flp plasmid. **C.** Ranked protein abundance in WT *E. coli* cells during exponential phase, as determined using quantitative mass spectrometry. NAPs (pink) are highlighted along with select other proteins. iBAQ: intensity-based absolute quantification. **D.** Read coverage at NAP loci in the genomes of ΔNAP1 to ΔNAP9 based on sequencing of genomic DNA and RNA. **E.** Viability of WT and ΔNAP9 cells sampled during different parts of the growth cycle (N=4 biological replicates each). Colony forming units (CFUs) per ml of culture were quantify by plating. Results are normalized by the OD_600_ of the cultures. **F.** Growth curves for WT, ΔNAP5 and ΔNAP9 in LB medium at 37°C. Characteristic doubling times and times to maximum growth rate, calculated from the growth curves (see Figure S3), are shown for WT, ΔNAP9, and all intermediate strains (N=3 biological replicates each, averaged over 4 technical replicates, ± SEM). **G.** Growth of the WT, ΔNAP5 and ΔNAP9 under different (stress) conditions (N=3 biological replicates each, averaged over 4 technical replicates, ± SEM). **H.** Representative STED images of WT and ΔNAP9 cells in exponential phase, along with quantification of relative cell and nucleoid dimensions. A total of 1,275 (1,331) WT and 1,925 (1,676) ΔNAP9 cells were analysed during exponential (stationary) phase. **I.** Cell length, cell width, and nucleocytoplasmic ratio of WT and ΔNAP9 cells in both exponential and stationary phase, as observed by confocal microscopy. **J.** Timeline showing the appearance and persistence of mutations during the construction of ΔNAP9.

We assumed that, even in rich medium conditions, bulk removal of all nine target NAPs would be lethal or strongly deleterious. We therefore opted for a serial deletion approach, leveraging existing single-gene deletion mutants from the KEIO collection (Baba et al. 2006), to remove one target NAP after another using P1 phage transduction (see Methods, Figure 1B), leaving behind only a 82 bp scar containing a 34 bp FRT site. Anticipating that, sooner or later, a fitness roadblock would emerge and prevent subsequent deletions, we reasoned that this step-wise approach would at least allow us to gradually edge up to the boundaries of what is possible and, by exploring multiple deletion paths (Figure S2), to circumnavigate epistatic traps (where the order of deletions blocked certain paths in an idiosyncratic, hard-to-predict manner). Ultimately, however, we persisted with only one path because the deletion journey – to our ongoing surprise – did not hit a sudden dead end: we deleted all nine target genes in succession.

Sequencing the genome of this strain, which we call ΔNAP9, and the genomes of all intermediate strains (ΔNAP1 to ΔNAP8, Table S1) using long-read Nanopore technology confirmed deletions of all NAP-encoding genes (Figure 1D). PCRs with primers targeting the inside of NAP genes (Table S2) further confirmed that the genes had not persisted outside their native genomic context. Deep sequencing of the transcriptome (described below) showed no RNA reads mapping to the NAP gene loci (Figure 1D), further confirming their effective deletion.

### No evidence for nucleoid decompaction in E. coli cells that lack all major NAPs

ΔNAP9 grows more slowly and has lower viability than its progenitor strain BW25113 (henceforth referred to as WT), with a doubling time approximately three times that of the WT strain (22 vs 63 min), and a notably extended lag phase (Figure 1E,F, Figure S3).

ΔNAP9 cells tend to be elongated and thinner compared to WT cells (Figure 1H,I) and often contain multiple nucleoids spread along the cell, pointing to defects in cell division and/or chromosome segregation. Nucleoids, as assessed by stimulated emission depletion (STED) microscopy, also appear thinner in ΔNAP9 (Figure 1H), reminiscent of nucleoids of *E. coli* cells treated with the beta-lactam antibiotic co-amoxiclav (Zagajewski et al. 2023). Interestingly, however, we find no evidence for nucleoid decompaction; the proportion of the total cell area occupied by the nucleoid, also known as nucleocytoplasmic (NC) ratio, is comparable to WT in both exponential and stationary phase (Figure 1I). This suggests that the absence of NAPs is effectively buffered in ΔNAP9 or, more heretically, that, compared to other factors such as molecular crowding or the charge-shielding effects of polycations, NAPs play a relatively minor role in chromosome compaction to begin with. This is arguably unexpected given prior observations from individual NAP deletion strains, e.g. Δ*dps* (Janissen et al. 2018), documenting measurable decompaction.

### Potential compensatory mutations along the path to ΔNAP9

Examining the growth patterns of intermediate strains (ΔNAP1-8), it is noticeable that growth performance does not decline monotonously with the number of NAPs removed (Figure 1F, Figure S3). In particular, we observe faster growth in ΔNAP5 compared to ΔNAP4. This could indicate that the removal of *ihfA* from the ΔNAP4 genetic background (Δ*hupA*Δ*hupB*Δ*hns*Δ*ihfB*) is, for unknown reasons, beneficial. Alternatively, and perhaps more likely, compensatory mutations might have arisen on the ΔNAP4 background prior to, during, or just after the construction of ΔNAP5. To explore this further, we sequenced the genomes of all intermediate strains using long-read sequencing (see Methods). We detect six mutations that emerged during the serial deletion process and became fixed along the way to ΔNAP9. Intriguingly, this includes mutations in DNA topoisomerase I (*topA*) and the replication initiator *dnaA* (Figure 1J, Table S3), which play key roles in supercoiling homeostasis and DNA replication, respectively, and are therefore prime candidates for compensatory evolution, as further discussed below. Notably, the mutation in TopA, causing a three amino acid deletion in its zinc-ribbon-like domain D8, which is in close proximity to single-stranded DNA during the topoisomerase cleavage cycle (Tan et al. 2015), is present in ΔNAP5 and its successor strains, but absent from ΔNAP4, suggestive of its involvement in the improved growth performance of ΔNAP5.

While ΔNAP9 grows slowly but robustly in rich medium, growth is markedly reduced under a variety of conditions that, for the WT strain, are only mildly stressful, including growth on minimal M9 medium (Figure 1G). Effects are particularly severe when cultures are treated with the DNA gyrase inhibitor novobiocin, suggesting that ΔNAP9 struggles to adaptively regulate supercoiling in the absence of its normal complement of NAPs, some of which – especially HU – are known to constrain negative supercoils (Broyles and Pettijohn 1986).

### The superhelical density of DNA in ΔNAP9 approaches known physiological limits

To measure the effects of NAP removal on supercoiling directly, we resolved DNA topoisomers of a pBR322 reporter plasmid introduced into WT, ΔNAP5 and ΔNAP9 cells using gel electrophoresis (Figure 2A,B, see Methods). Plasmid DNA in WT *E. coli* is mildly negatively supercoiled, with an average superhelical density of −0.033. In comparison, DNA topology in ΔNAP5 and ΔNAP9 is more relaxed, their superhelical density 10% lower than WT levels (−0.030), close to a previously proposed threshold beyond which *E. coli* cells struggle to survive (Champion and Higgins 2007; Rovinskiy et al. 2019). Assuming that the mutation in TopA reduces its relaxation activity, this change is clearly insufficient to restore WT levels of negative supercoiling. These observations are consistent with a model where ΔNAP5 and ΔNAP9 have a reduced capacity to constrain negative supercoils and becomes more sensitive to novobiocin-mediated inhibition of DNA gyrase, which now has a heavier burden to shoulder in maintaining physiologically necessary levels of negative supercoiling across the genome.

**Figure 2.**
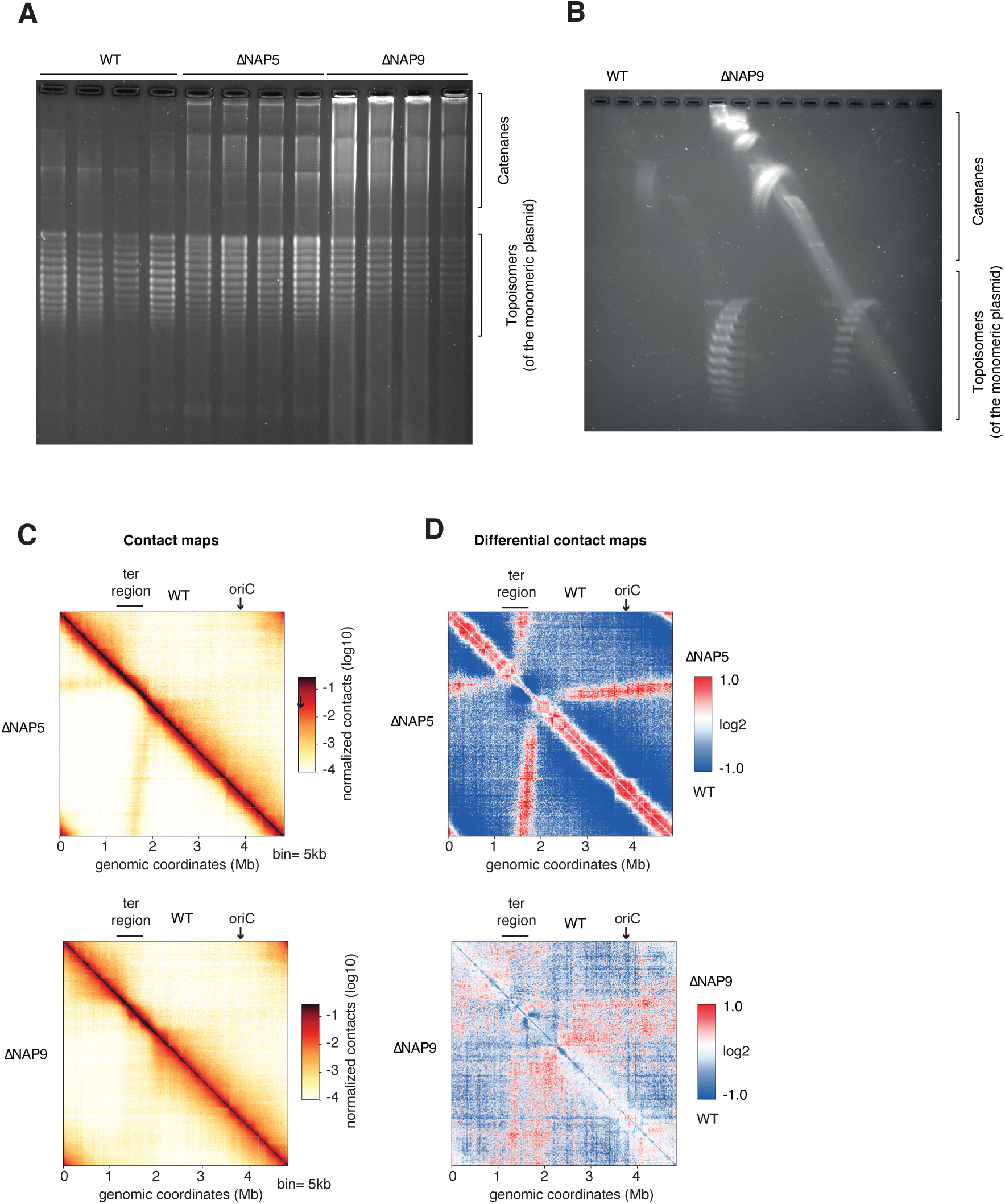
Effects of NAP removal on supercoiling and chromosome conformation. **A.** 1D electrophoresis gel showing the distribution of topoisomers of the pBR322 plasmid extracted from WT, ΔNAP5 and ΔNAP9 strains in exponential phase. Four biological replicates per strain are presented. Higher molecular weight species, likely catenanes, are visible at the top of the gel. **B.** 2D electrophoresis gel showing the distribution of topoisomers of the pBR322 plasmid extracted from WT or ΔNAP9 strains in exponential phase. Plasmids are resolved by their degree of DNA supercoiling in the first dimension, and by the sense of the supercoiling in the second dimension. Chloroquine is used as a DNA intercalating agent in both 1D and 2D gels to relax the excess of negative supercoiling and help visualising the topoisomers distribution. **C,D.** Contact maps (C) and ratio maps/differential contact maps (D) for WT, ΔNAP5, and ΔNAP9, illustrating the chromosomal contact intensities across the *E. coli* genome.

Beyond effects on superhelical density, the gels also reveal a pronounced tendency for plasmids in ΔNAP5 and ΔNAP9 to form larger species (Figure 2A,B), which likely correspond to catenanes (multimers that arise from interlinked monomers).

### Higher-order chromatin structure is largely undisturbed in ΔNAP9

To further explore the impact of the absence of nine NAPs on DNA topology and nucleoid architecture, we performed Hi-C experiments on ΔNAP9, ΔNAP5, and WT during exponential growth (see Methods). The WT contact map (Figure 2C) is consistent with prior maps in *E. coli* K-12, showing a constrained, contact-dense terminus region, flanked by longer-range interaction domains (Lioy et al. 2018). The contact map for ΔNAP5 exhibits a strong enrichment for short- and medium-range contacts, illustrated by the red diagonal in the ΔNAP5/WT ratio map (Figure 2D) and a concomitant depletion in longer-range contacts, as previously observed for a Δ*hupA*Δ*hupB* strain (Lioy et al. 2018). The most striking feature of the ΔNAP5 contact map is the emergence of a cross-shaped signal, indicating enriched contacts between the regions adjacent to the terminus macrodomain and the rest of the genome (up to and including the origin region). This pattern is strongly reminiscent of the “butterfly wing” pattern previously observed when inactivating topoisomerase IV using thermosensitive alleles of either the parC or parE subunit (Conin et al. 2022). In that instance, the butterfly wing pattern was attributed to a decatenation defect (i.e. the failure to disentangle sister chromatids following replication), as linearization of the *E. coli* chromosome strongly attenuated the signal (Conin et al. 2022). We suggest that ΔNAP5 suffers from a similar defect, i.e. a compromised capacity to resolve interlinked chromosomes and segregate them in a timely and coordinated way.

In comparison, the contact map of ΔNAP9 appears much more similar to WT (Figure 2C,D). The butterfly wing pattern is strongly attenuated (and less narrowly localized) and there is a decrease in short-range contacts in ΔNAP9 that appears relatively homogeneous and is evident along the entire genome, resulting in a thin blue stripe along the diagonal of the contact ratio map (Figure 2D). Recent results suggest that this might reflect reduced supercoiling at the level of promoters, which have been shown to constrain contacts towards more local interactions (Bignaud et al. 2024). Considering contact frequency as a function of genomic distance (contact decay, Figure S4) further emphasizes the relative similarity of overall nucleoid architecture between ΔNAP9 and WT, a result that is consistent with our imaging results and conservation of the NC ratio reported above, but unexpected given impacts previously seen in Hi-C maps following the removal of individual NAPs (Lioy et al. 2018; Conin et al. 2022; Gavrilov et al. 2025) and the stronger accumulation of catenanes for the reporter plasmid in ΔNAP9, which might have led one to predict a stronger butterfly wing pattern in ΔNAP9 versus ΔNAP5.

### Global changes in the ΔNAP9 transcriptome are predicted by wildtype expression level

Through a variety of mechanisms, from direct occlusion of promoters to affecting RNA polymerase loading and progression, NAPs are intimately involved in the regulation of transcription (Ge et al. 2025; Amemiya et al. 2021b; Dillon and Dorman 2010). To explore how bulk NAP removal impacts transcription, we surveyed transcriptional responses in WT, ΔNAP5, and ΔNAP9 cells using RNA-Seq.

The transcriptome of the deletion strains is clearly distinct from that of WT cells (Figure 3A). Dysregulation is global, with 3424 out of 4337 (79%) genes significantly up- or down-regulated when comparing WT and ΔNAP9 exponential phase transcriptomes (Figure 3B), with a comparable number of genes (3569/4337=82%) affected in stationary phase. Similar numbers of genes are up-versus down-regulated, but large (>2-fold) changes are more prominent amongst upregulated genes (Figure 3B).

**Figure 3.**
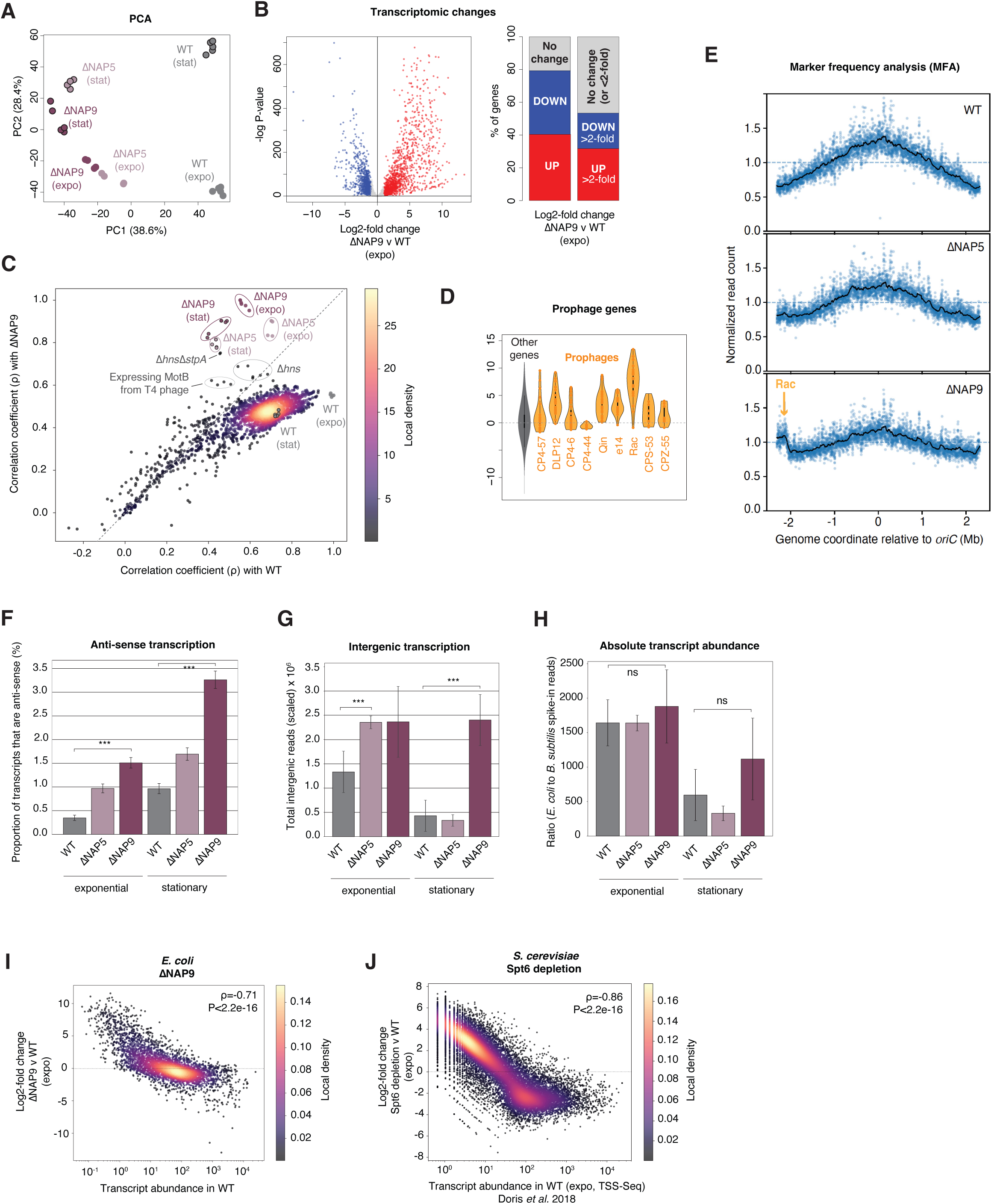
Effects of NAP removal on transcription. **A.** Principal component analysis (PCA) of RNA-Seq profiles for WT, ΔNAP5 and ΔNAP9 in exponential and stationary phase. **B.** Differential expression of genes in ΔNAP9 versus WT. Genes significantly up- or down-regulated are coloured in red and blue, respectively. **C.** Similarity of transcription profiles in WT, ΔNAP5 and ΔNAP9 compared to a curated collection of transcriptomic datasets (see main text), using one replicate of WT (expo) and one replicate of ΔNAP9 (expo) as reference points. Conditions of particular interest are highlighted; all correlation coefficients are provided in Table S4. **D.** Upregulation of genes associated with cryptic prophages in ΔNAP9. Differential regulation of genes outside of prophage regions are provided for comparison **E.** Marker frequency analysis (MFA, see Methods) based on exponentially growing WT, ΔNAP5, and ΔNAP9 cultures. The terminus-adjacent location of the prophage Rac, which is associated with higher than expected coverage, is highlighted. Read counts are normalized in each strain. **F.** Proportion of sequencing reads mapping anti-sense to known transcription units in WT, ΔNAP5 and ΔNAP9. Error bars indicate SEM. **G.** Sequencing reads in WT, ΔNAP5 and ΔNAP9 mapping to intergenic locations, irrespective of their orientation relative to adjacent genes (also see Figure S5). **H.** Comparison of absolute mRNA abundance across samples relative to spike-in controls, obtained by addition of a fixed quantity of *B. subtilis* cells. **I.** Changes in transcript abundance in ΔNAP9 as a function of WT expression levels, illustrating genome-wide homogenization of the transcriptional landscape in ΔNAP9. WT transcript abundance (transcripts per million, TPM) is provided as the mean across the three most correlated WT replicates. ***J.*** Changes in transcript abundance following Spt6 depletion in *S. cerevisiae* as a function of WT expression levels. Transcript counts were determined using transcription start site sequencing (TSS-Seq; see Doris *et al*. 2018). ***P<0.001 Wilcoxon rank-sum test.

To understand how unusual these changes are in the context of *E. coli* physiological flexibility, we integrated our RNA-Seq data with a compendium of 2207 previously compiled and consistently re-processed *E. coli* MG1655 transcriptomic datasets (see Methods), which cover a large variety of drug treatments, genetic changes, and growth on different carbon sources (Tjaden 2023). A large majority (92%) of previously measured transcriptomes are more similar to our WT transcriptome than they are to the ΔNAP9/ΔNAP5 transcriptomes (Figure 3C, Table S4). The datasets where we do find genuinely greater affiliation to the ΔNAP9/ΔNAP5 compared to the WT transcriptome, share a strong common theme, coming from strains deleted for *hns,* alone or in combination with its paralog *stpA,* or expressing the T4 phage protein MotB, which has been suggested to interfere with H-NS binding to DNA (Patterson-West et al. 2018). Perhaps unsurprisingly, genes that are de-repressed in a Δ*hns*Δ*stpA* double deletion background are similarly de-repressed in ΔNAP9. However, transcriptomic changes in Δ*hns*Δ*stpA* still only very partially capture the changes observed in ΔNAP9 (ρ=0.35, Figure S5).

Consistent with the deletion of *hns* and *stpA*, and their known roles as xenogenic silencers, we find that genes belonging to the cryptic prophages present in BW25113 (Figure 1A) are consistently upregulated in ΔNAP9 (Figure 3D). This is particularly pronounced for the prophage Rac, which is a known direct target of H-NS (Hong et al. 2010). Marker frequency analysis (MFA, see Methods) suggests that Rac actively replicates in ΔNAP9 (Figure 3E).

Looking beyond annotated protein-coding genes, we also observe an increase in non-canonical transcription events: antisense transcription, where the transcript is produced from the strand opposite to a known transcription unit, increases more than 3-fold in ΔNAP9 (Figure 3F, Figure S5) and a ∼2-fold higher proportion of reads maps to intergenic DNA (Figure 3G), pointing to a global loosening of transcriptional control.

In eukaryotes, histone deletion or depletion leads to a more promiscuous transcriptional landscape, characterized by the activation of previously cryptic intra- and intergenic promoters (Gossett and Lieb 2012; Wyrick et al. 1999; Doris et al. 2018; Kaplan 2003; Chari et al. 2019). This can, in principle, be linked to greater transcriptional activity in absolute terms, i.e. more transcripts being generated per cell, as histone removal renders a previously restrictive ground state more permissive. Alternatively, chromatin might not be rate limiting for transcription and absolute transcriptional output remains the same, but, following unmasking of previously cryptic sites, RNA polymerases are titrated away to newly accessible promoters leading to a reduction of transcriptional activity at the originally active loci. This effect has previously been reported for an *E. coli hns* deletion strain (Lamberte et al. 2017).

Approximating the total transcriptional output per cell using spike-ins (see Methods), we find no difference between WT and ΔNAP9 cells (Figure 3H), suggesting that bacterial chromatin is not rate-limiting for total transcriptional output. Instead, the pervasive transcriptomic changes we observe are consistent with global reallocation of transcriptional activity, which is happening in a strikingly predictable way: The most highly expressed genes in the WT are most strongly downregulated in ΔNAP9 whereas the most lowly expressed genes in the WT experience the strongest upregulation, resulting in an overall homogenization of transcription across the genome (Figure 3I).

A tendency for genome-wide homogenization of transcriptional output has previously been observed for other proteins with a global role in genome regulation, for example following the deletion of reverse gyrase in the archaeon *Thermococcus kodakarensis* (Villain et al. 2025), the inhibition of topoisomerase I in the bacterium *Dickeya dadantii* (Pineau et al. 2022), novobiocin-mediated inhibition of DNA gyrase in *E. coli* (Lamoureux et al. 2023), and the depletion of cohesin from mouse thymocytes (Seitan et al. 2013) (Figure S6). In all these cases, however, the relationship between WT expression and transcriptional change following the intervention is much less pronounced (|ρ|<0.22 in all cases) than what we observe here (ρ=-0.71). The condition that most closely resembles our results, especially when taking cryptic promoters into account, is what happens to nascent transcription in budding yeast upon depletion of Spt6, a histone chaperone critical for reassembling nucleosomes following the passage of RNA polymerase (Doris et al. 2018; Cheung et al. 2008; Kaplan 2003). Here too, the up- and down-regulation of individual loci is well predicted by wildtype transcription levels (Figure 3J, Figure S6), suggesting hitherto unappreciated similarities in how chromatin structures genome-wide transcriptional output in prokaryotes and eukaryotes.

To establish how bulk NAP removal affects the proteome, we measured protein levels in WT and ΔNAP9 using quantitative mass spectrometry (see Methods). Overall, changes in transcript and protein abundance are well-correlated for both WT and ΔNAP9 (Figure 4A).

**Figure 4.**
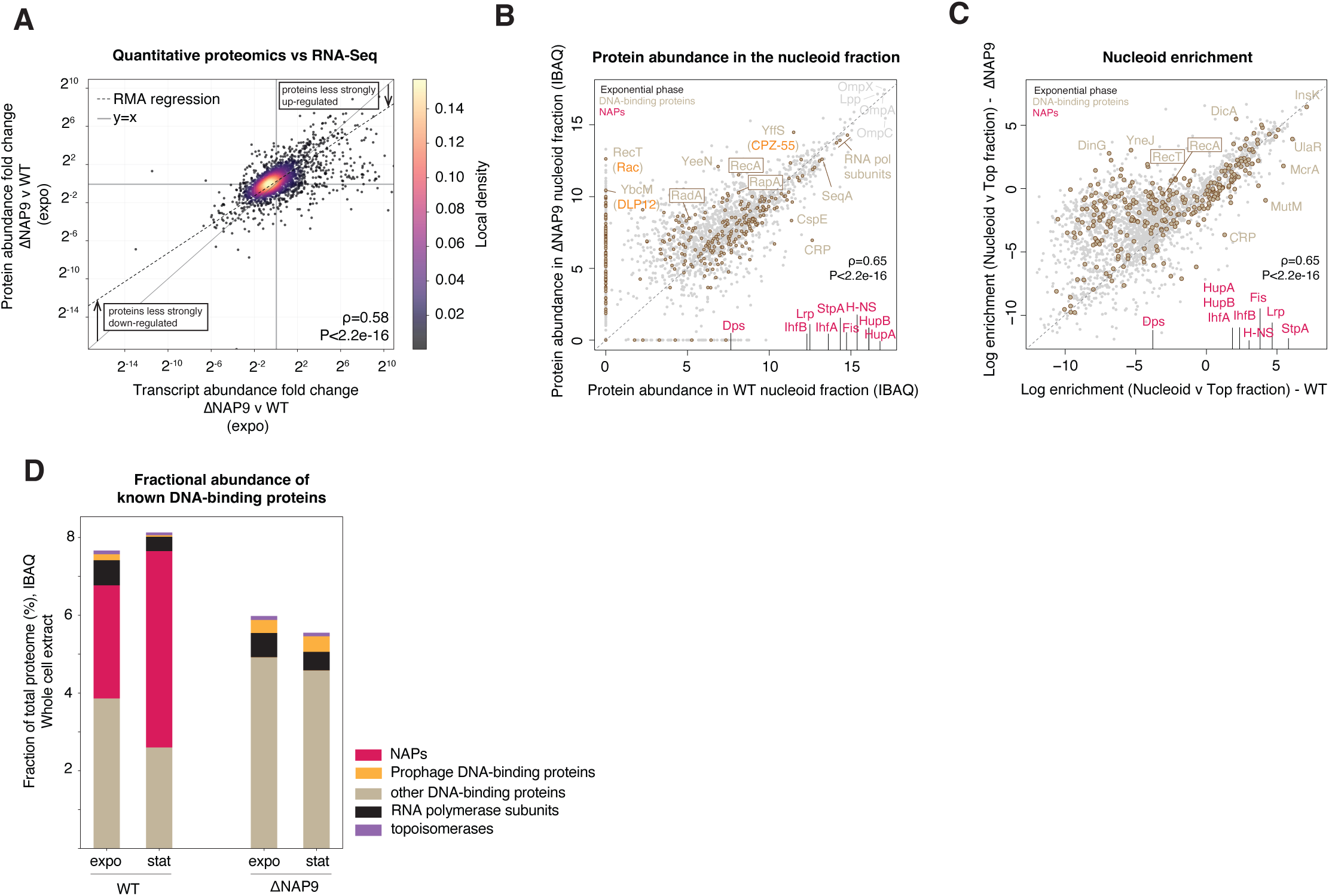
Effects of NAP removal on the cellular and nucleoid-associated proteome. **A.** The magnitude of change in protein abundance in ΔNAP9 compared to WT is smaller than the magnitude of change for the corresponding transcripts, as evident in the location/slope of the Reduced Major Axis (RMA) regression line, which describes the empirical relationship between transcript and protein changes, compared to the x = y line **B,C.** Abundance (B) and relative enrichment (C) of proteins in the nucleoid fraction (compared to the top fraction, see main text) comparing WT and ΔNAP9 exponential phase cells. The behaviour of DNA-binding proteins is highlighted (ivory with brown outline). The abundance of NAPs (pink) in the WT is indicated along the x-axis. D. Fraction of the total proteome invested in producing different types of DNA-binding proteins.

There is some range compression (i.e. protein abundance changes are less pronounced than changes in the abundance of the corresponding RNAs, for both up- and down-regulation), but homogenization of the transcriptome clearly feeds through to the proteome.

### Limited remodelling of the nucleoid proteome upon bulk NAP removal

Although ΔNAP9 lacks the main building blocks of *E. coli* chromatin, this does not mean, of course, that the chromosome is suddenly devoid of proteins. Numerous proteins will still be associated with DNA, including processive proteins such as DNA/RNA polymerases and a large collection of sequence-specific transcription factors. But does NAP removal precipitate changes to nucleoid composition that might hint at mechanisms by which the system compensates for the absence of its native NAPs? For example, do we observe upregulation or increased chromatin association of certain transcription factors that might step in to fill the functional gap left by the deletion of *E. coli’s* regular NAP complement? To investigate this, we again quantified protein abundance via quantitative mass spectrometry but this time specifically for nucleoid-enriched subcellular fractions obtained via sucrose gradient centrifugation (Ohniwa et al. 2011; Hocher et al. 2022) (see Methods).

Consistent with prior results (Ohniwa et al. 2011), we find our target NAPs to be highly abundant in the nucleoid fraction of WT cells and highly enriched compared to the top fraction, which predominantly contains cytosolic proteins (Figure 4B). Note here that, as previously discussed (Ohniwa et al. 2011; Hocher et al. 2022), the nucleoid enrichment protocol does not isolate the nucleoid exclusively but also contains numerous proteins associated with the membrane, presumably because these two entities are physically linked via transertion.

We then searched for deviations in protein abundance and enrichment between WT and ΔNAP9 samples. On a background of comparative stasis, three things stand out: First, perhaps the most obvious candidate proteins to act as next-in-line stand-ins for the regular set of NAPs, proteins such as CspE, CRP, SeqA or Hfq, show no conspicuous increase in abundance or enrichment in ΔNAP9 nucleoids (Figure 4B,C). At the same time, non-NAP DNA-binding proteins as a group jointly make up a larger fraction of the total proteome than they do in the wildtype (Figure 4D). The nucleoid is thus partially re-populated by many different DNA-binding proteins that individually experience small increases in abundance.

Second, of the known DNA-binding proteins whose functional roles have been studied, RecA is noticeably more abundant and more enriched in the ΔNAP9 nucleoid, suggesting greater recruitment of the cellular pool of RecA to the chromosome (Figure 4B). Alongside RecA, two of its known interaction partners are also more abundant in the nucleoid: RapA and RadA. RapA is involved (alongside TopA) in the resolution of R-loops, which can arise during transcription-replication conflicts, perhaps suggesting that, in the absence of NAPs, such conflicts become more common. RadA, on the other hand, participates alongside RecA in homologous recombination, including in the context of double strand break repair, suggesting that genome integrity might be affected in ΔNAP9. Third, some of the proteins for which abundance in the nucleoid fraction has changed most dramatically are derived from prophages, illustrated in particular by YffS, YeeN, and RecT (Figure 4B). This is perhaps not surprising given the global upregulation of prophages we reported above (Figure 3D), but begs the question whether prophage-encoded DNA-binding proteins might, inadvertently, act to compensate for the loss of *E. coli’s* regular complement of NAPs, especially in light of prior work that suggested that activation of cryptic prophages in *E. coli* BW25113 can be beneficial to the host by increasing its ability to tolerate a variety of stressors (Wang et al. 2010). The next set of experiments, rather by accident, suggests that, in fact, the opposite is true.

### Repeated loss of prophages during experimental evolution of ΔNAP9

We highlighted above that a small number of mutations fixed during the construction of ΔNAP9. It seems likely that at least some of these mutations help compensate for the loss of NAPs, although their precise contributions remain to be determined (see below). At the same time, it also seems likely that capacity exists for further compensatory evolution. To test this, we subjected five replicate cultures of ΔNAP9 (alongside five replicate WT cultures as controls) to 50 days (∼ 500 generations) of experimental evolution, sub-culturing daily under benign conditions to facilitate the fixation of mutations that would increase the fitness of these strains (Figure 5A, see Methods). At the end of the experiment, we compared growth rates between the ΔNAP9 ancestor and the evolved strains, finding much improved growth rates across the board (Figure S7). To measure fitness recovery directly, we carried out pairwise competitive fitness assays. All evolved lines rapidly outcompeted their ΔNAP9 ancestor (Figure 5B). We then sequenced the evolved lines and looked for genetic changes that might underpin fitness recovery. Genomes of two of the lines (ΔNAP9_ev2, and ΔNAP9_ev3, see Figure 5C) carry the exact same set of mutations, perhaps indicative of cross-contamination, but otherwise mutation patterns are unique, supporting independent events across the different lines. Strikingly, every single one of the evolved lines carries large deletions in parts of the genome that harbour prophages, specifically Rac, DLP12, CPZ-55, and CP4-6 (Figure 5C,D). The largest deletion, spanning Rac, encompasses 145 genes (174.9kb). In the case of DLP12, the genomic deletion maps neatly onto the phage’s attP sites, indicative of mobilization, whereas deletion boundaries for the Rac and CPZ-55 are more heterogeneous, suggesting a mixture of specific mobilization and genomic deletions. In one line (ΔNAP9_ev4), a large region downstream of Rac has been deleted whereas the core Rac region itself remains intact. This region contains numerous genes strongly upregulated in ΔNAP9 and is part of the same H-NS-repressed genomic domain as Rac itself (Figure S8).

**Figure 5.**
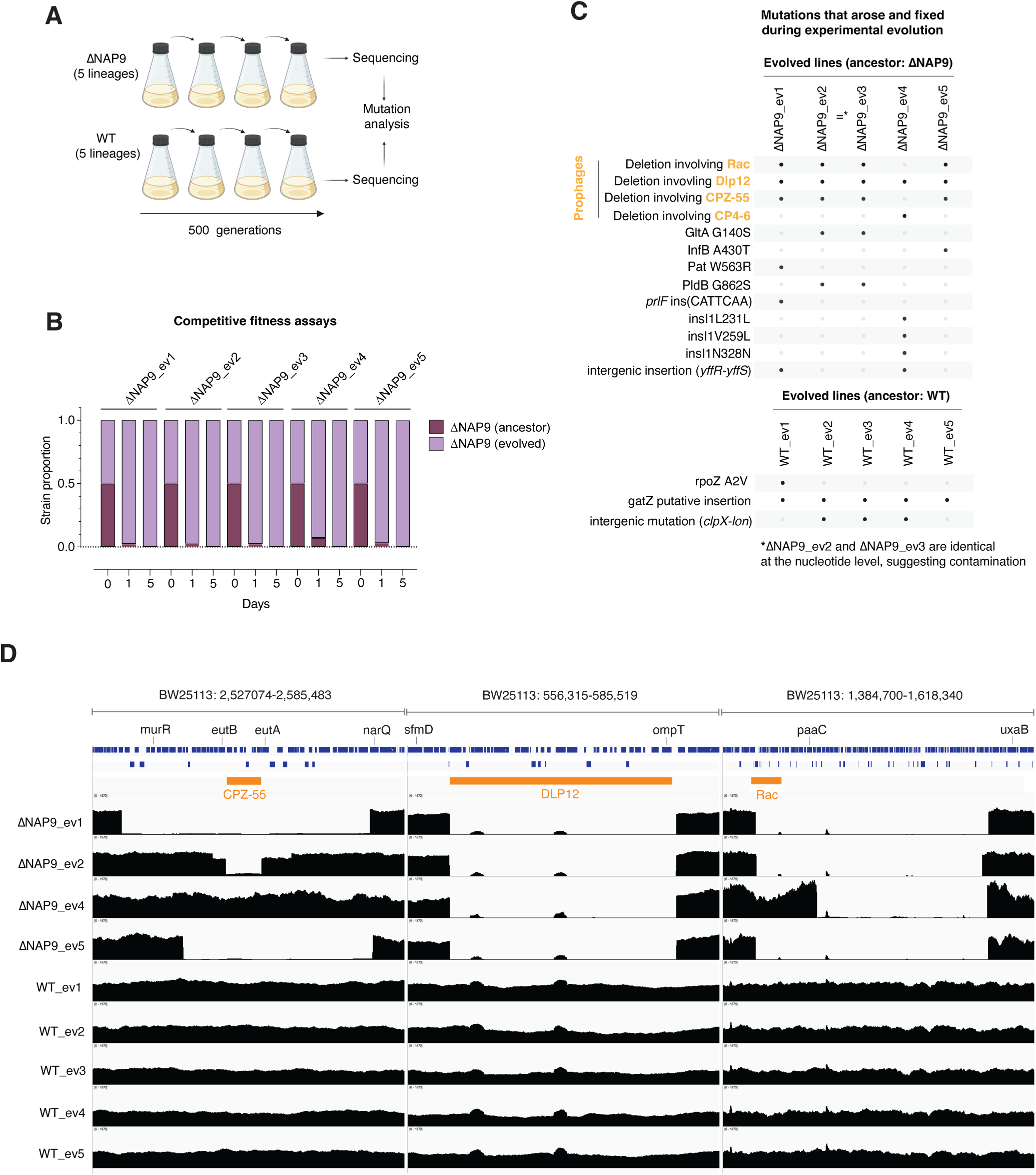
Experimental evolution of ΔNAP9. **A.** Schematic of experimental evolution set-up, where five independent lineages of ΔNAP9 and the WT were sub-cultured for 500 generations in LB medium. Genomes of the evolved lineages were sequenced to identify genetic changes that occurred during evolution. **B.** Pairwise competitive fitness assays between the ΔNAP9 ancestor and each evolved lineage. The ancestral strain is rapidly outcompeted in all cases. **C.** Summary of mutations that fixed in different lineages evolved from the ΔNAP9 (or WT) ancestor. **D.** Coverage traces at genomic loci affected by deletions of (or nearby) cryptic prophages CPZ-55, DLP12, and Rac.

These results suggest that activation of cryptic prophages and co-repressed genomic neighbourhoods, rather than helping to compensate for the loss of NAPs by donating DNA-binding proteins with potentially useful properties for nucleoid function, induces a considerable fitness burden.

## DISCUSSION

The serial deletion of nine NAPs from *E. coli* demonstrates that it is possible for a bacterial cell to carry out basic cellular functions - to grow, replicate, and divide – in the absence of its native complement of chromatin proteins. This is unexpected. Prior work described even a simple *ΔhupA/ΔhupB* double mutant as “extremely compromised” (Huisman et al. 1989), genetically unstable (Ogawa et al. 1989), and subject to rapid acquisition of suppressor mutations (Malik et al. 1996), and previous efforts to construct higher-order NAP deletions had failed (Yasuzawa et al. 1992). This suggests that the genesis of ΔNAP9 involved at least some serendipity, be that in the fortuitous choice of an amenable genetic background or the emergence of compensatory mutations that allowed the removal of additional NAPs.

The existence of ΔNAP9 does not imply, of course, that chromatin is unimportant for fitness. Our findings reinforce known parallels between prokaryotes and eukaryotes regarding the adaptive role chromatin plays in keeping in check selfish elements such as (pro)phages, retroviruses, and transposable elements. And although the NAPs deleted here may not be required for cellular life per se, the ability of *E. coli* to respond adaptively to changing conditions is very much compromised (Figure 1G), consistent with a known involvement of NAPs in mediating physiological responses to environmental change (Hołówka and Zakrzewska-Czerwińska 2020; Ge et al. 2025). Genes continue to be expressed at levels compatible with fundamental growth requirements, but one of the mechanisms by which the cell can dynamically reshape how genomic information is selectively deployed, is lost. One striking manifestation of this is the homogenization of transcriptional output, presumably linked to global re-allocation of RNA polymerase activity (Lamberte et al. 2017), mirroring what happens in yeast when histone deposition onto DNA is disrupted (Doris et al. 2018). In both eukaryotes and bacteria, without chromatin as a lithe adaptive stencil, genomic activity, and the transcriptional image that emerges from it, lack definition.

Our findings are consistent with – and extend – recent work showing that the effect of transcription factor binding on gene expression in *E. coli* can be predicted by simple thermodynamic models. Using a combination of theory and experiments, Brewster and colleagues showed that, in a regime where transcription factor binding facilitates recruitment of RNA polymerase to the promoter or stabilizes the association, the expected change in transcription output scales inversely, in sign and magnitude, with baseline levels of gene expression (Parisutham et al. 2025; Guharajan et al. 2025). In other words, the most highly expressed genes tend to be the most downregulated and the most lowly expressed genes the most upregulated. This is what we observe here, but happening on a genome-wide scale upon removal rather than addition of DNA-binding proteins. The latter is consistent with the ubiquity of NAPs in the vicinity of promoters, the former with a model where, on average, the presence of NAPs prevents or destabilizes interactions between promoters and the core transcription machinery.

### Limitations

As noted above, sequencing of ΔNAP9 revealed a number of mutations that arose and fixed in the process of building ΔNAP9, including in genes with known involvement in nucleoid biology, notably *dnaA* and *topA*. These mutations are exciting because – if they do indeed act in a compensatory manner - they provide a window onto the genetic and evolutionary pathways that mitigate fitness loss and through which chromatin loss becomes possible. By implication, they might also tell us which of the many processes in which chromatin is involved is most acutely compromised. On the flipside, the presence of these mutations complicates our evaluation of bulk chromatin loss. In an ideal scenario, WT and ΔNAP9 would be isogenic except for the deletions we introduce. Clearly, this is not the case here and the genomic consequences we assay are a joint consequence of the deletions themselves and the mutations that accompany them. Given the unexpectedly high viability of ΔNAP9, we strongly suspect that at least some of the mutations do play a compensatory role. The mutations in *dnaA* and *topA* are prime candidates here given the known (and on occasion joint) roles of these proteins in DNA transactions. For example, DnaA and TopA are involved alongside IHF/HU in the initiation of replication at oriC (Magnan and Bates 2015), so it is tempting to speculate that the removal of HU (and IHF) proteins selects for compensatory mutations in these proteins. Mutations in *rpsB,* which encodes a 30S ribosomal subunit, and *mnmE,* involved in tRNA modification, could also contribute to compensation, although effects here are harder to rationalize with reference to known chromatin biology.

But whether and how precisely any of the mutations in ΔNAP9 contribute to its survival remains to be established, including how specifically compensatory effects are tied to chromatin biology. Mutations in *topA* are common in experimental evolution studies of *E. coli* (Woods et al. 2011; Bachar et al. 2019; Crozat et al. 2005, 2010), including under conditions where the adaptive challenge is not directly linked to chromatin. Since TopA, DNA gyrase, and similar global regulators of supercoiling affect many genes simultaneously, mutations in these genes might be considered an evolutionary Hail Mary – causing multiple changes in expression, one of which has an outside chance of getting the maladapted organism out of its bind. It will be instructive, in the future, to attempt to revert the mutations in ΔNAP9 to isolate their specific contributions to fitness and thereby further delineate the limits to life without chromatin.

## Conclusion

Whatever the precise combination of factors that facilitated its construction, the fact that ΔNAP9 exists shows that there are evolutionary pathways that enable bulk removal of chromatin proteins. It is another example, if one was needed, of the phenomenal plasticity of prokaryotic systems, their ability to cope with and ultimately adapt to radical changes in their cellular set-up, and illustrates the potential for fundamental re-engineering of prokaryotic genomes in general and prokaryotic chromatin in particular. To understand the practical scope for chromatin engineering, it will be important in the future to establish the precise genetic and physiological conditions that enable chromatin removal/depletion, as this will inform efforts to carry out similar experiments in different genetic backgrounds or even different species, including those where individual NAPs are essential, as is the case for HU in *B. subtilis* (Karaboja and Wang 2022). In this context, it would be interesting to repeat the serial deletion experiment, including using a different deletion order, to understand whether the genomic adaptations that emerge are conditioned by the specific deletion history of the strain.

Our demonstration that chromatin proteins can be deleted *en masse* might also open up new opportunities not only for studying the broader evolutionary drivers of chromatinization but also for understanding the function of individual chromatin proteins. Historically, the unique contributions of individual NAPs *in vivo* and how they interact to jointly shape genome regulation has been difficult to dissect because of widespread redundancy. Strains like ΔNAP9 offer the chance to introduce individual NAPs, alone or in combination, in a controlled manner to study their *in vivo* behaviour free from interference by other NAPs.

## METHODS

### Strain Construction by phage transduction

NAPs deletion mutants were constructed by sequential transduction of individual deletions from the Keio collection in *E. coli* BW25113 (Baba et al. 2006). The IS1-free virulent P1 bacteriophage (P1*vir*) used for the transduction (Fehér et al. 2012) was kindly provided by Tamas Fehér. Preparation of P1*vir* lysates and transduction were performed as described by (Thomason et al. 2007) with minor modifications. An overnight culture of the recipient strain was grown at 37°C with shaking in LB medium. About 4 x 10^9^ cells were collected by centrifugation for 30 s at 11,000 x g. Cells were resuspended in 250 µl of P1 salts (5 mM CaCl_2_ and 10 mM MgSO_4_, filtered 0.2 µm) and P1*vir* lysates were added to reach a multiplicity of infection of ∼1. Cells with P1*vir* were incubated for 25 min at 37°C to get a single round of infection, then the infection was stopped by addition of 200 µl of 1 M sodium citrate and 1 ml of LB. Cells were incubated for 90 min at 37°C with shaking for recovery, before being harvested by centrifugation for 30 s at 11,000 x g and plated on LBKC plates (LB plates supplemented with 30 µg/ml of kanamycin and 5 mM of sodium citrate).

Depending on the growth rate of the transductants, 8 colonies were screened by PCR after 20 to 48 hours using primers external to the targeted genes, and then re-streaked twice on LBKC plates to eliminate any remaining P1*vir* particles.

Two individual clones were then transformed by standard electroporation with pBAD-Flp (Scholz et al. 2019) . Briefly, 1 to 2 ml of overnight culture were pelleted for 30 s at 11,000 x g, washed on ice 5 times with ice-cold 10% glycerol, resuspended in 100 µl with 1 to 100 ng of plasmid. Cells were electroporated in 0.1 cm gap cuvette with a BioRad Gene Pulser II electroporator (25 µF, 200 Ω, 1.7 kV) before incubation for 90 to 120 min in 1 ml LB at 30°C with shaking. After the recovery, cells were plated on LB supplemented with 33 µg/ml of chloramphenicol and incubated for up to 48 h at 30°C.

After deleting *hupA*, *hupB*, *hns*, *ihfB*, and *ihfA* we did not manage to transform the subsequent strains with pBAD-Flp to recycle the kanR cassette. This suggested a dependence of the plasmid on one of the deleted NAPs. Three putative IHF binding sites were identified in the sequence of pBAD-FLP using FIMO (Grant et al. 2011), strengthening this hypothesis. We therefore decided to clone *ihfA* and *ihfB* on pBAD-Flp, under the control of a rhamnose-inducible promoter to allow propagation of pBAD-Flp in the strains deleted for *ihfA* and *ihfB*. This pBAD-Flp-ihf plasmid was used for all the daughter strains of ΔNAP4. KanR cassette excision was performed by culturing the mutants transformed with pBAD-Flp overnight at 30°C, in LB supplemented 33 µg/ml of chloramphenicol, and 1% of L-arabinose to induce the flippase gene expression. For strains transformed with pBAD-Flp-ihf, 0.2% of rhamnose was added in addition to enable plasmid maintenance.

Following kanR cassette excision, pBAD-Flp and pBAD-Flp-ihf plasmids were cured from mutant strains by culturing them above 37°C. Briefly, cultures were diluted at 1/1000 in LB and incubated for a few generations at 37°C before being plated on LB and incubated overnight at 37°C. For the late deletions mutants (ΔNAP6+), the pBAD-Flp-ihf plasmid was harder to lose and 42°C instead of 37°C was used to cure the plasmid.

### Confirmation of deletions

Individual mutant clones were then phenotypically screened on LB plates supplemented either with kanamycin (no growth expected after cassette excision) or chloramphenicol (no growth expected after curation of pBAD-Flp or pBAD-Flp-ihf). Clones harboring expected phenotypes were screened by PCR with primers external and internal to the relevant NAP genes, and primers internal to kanR cassette (Table S1). Genomes of selected clones that had passed screening were sequenced using the Oxford Nanopore technology (ONT) bacterial whole genome sequencing service from Plasmidsaurus.

### Long-term evolution experiment

Five culture tubes containing 5 ml LB were inoculated with the ΔNAP9 strain. These cultures were propagated for about 500 generations (50 subcultures) by serial 1/1000 dilutions in 5 ml of LB, at 37°C, 180 rpm in a dedicated INFORS Ecotron shaking incubator. As a control, 5 WT lineages were cultured in the same way starting from the WT BW25113 strain. Genomic DNA of the evolved lineages was extracted using the Monarch Spin gDNA Extraction kit (New England Biolabs) and sequenced using the ONT whole bacterial genomes sequencing service from Plasmidsaurus.

### Mutation calling

ONT reads were mapped onto the BW25113 reference genome (NZ_CP064677.1) using Minimap2 (Li 2018). The resulting sam files were converted to bam files, sorted and indexed using SAMtools (Li et al. 2009). Mutation calling was performed using Clair3 (Zheng et al. 2022) with the --haploid_precise and --enable_long_indel options for the WT and ΔNAP1-9, and the --haploid_sensitive and --enable_long_indel options for the evolved strains (see GitHub repository at https://github.com/PaulVillain/NoNAPs_project for details). With the haploid_precise parameter, all the called mutations were considered, while with the haploid_sensitive parameter, only the mutation with an allele frequency above 0.4 were considered. All putative mutations were inspected on IGV for validation.

### Competition experiments

The fitness of the evolved ΔNAP9 lineages was assessed against the initial ΔNAP9 strain by via pairwise competition experiments. Independent co-cultures of the initial ΔNAP9 strain and the individual evolved lineages were conducted for ∼50 generations, at 37°C and 200 rpm, in 6 ml of LB, through 5 rounds of overnight subcultures after 1/1000 dilution in LB. For each co-culture, three biological replicates were inoculated from pre-cultures, at an equal OD_600_ for the initial ΔNAP9 strain and the relevant evolved ΔNAP9 lineage. Genomic DNA was extracted from the co-cultures with the Monarch gDNA extraction kit (NEB), just after mixing the initial pre-cultures (T0), after the first culture (T1) and after 5 subcultures (T5). The relative proportions of the initial ΔNAP9 strain and the ΔNAP9 evolved lineages in the co-cultures were then estimated by qPCR using the Luna Universal Master Mix (NEB). Primers located in a portion of the genome common to the initial ΔNAP9 strain and the evolved lineages were used to quantify the total cell population. Primers located in a deletion close to the Rac prophage common to all evolved lineages were used to quantify the ΔNAP9 initial strain in the co-culture. The qPCR reactions were set up with 40 pg of gDNA and run in a Rotor-Gene Q thermocycler (Qiagen) with the following program: 95°C 1 min, (95°C, 15s; 60°C, 30s) 40 cycles. The qPCR results were analysed using the Rotor-Gene Q Series software. The relative proportion of the initial ΔNAP9 strain and the evolved lineages was extrapolated from the 2^-ΔCt^.

### Transcriptomics

Frozen pellets of exponential phase (OD_600_ = 0.4) or stationary phase (OD_600_ = 2 to 8) cultures of WT, ΔNAP5 and ΔNAP9 were mixed with a fixed amount (0.015 OD_600_ units) of a *Bacillus subtilis* 168 culture as a spike-in. Pellets were then processed with the RNeasy kit (Qiagen), following the standard bacteria protocol, including the extra DNase step. Cells were lysed using 425-600 µm acid-washed glass beads (Sigma-Aldrich) and a TissueLyser (Qiagen). Quality and concentration of total RNA samples was assessed on agarose gels and then with a Bioanalyser 2100 (Agilent) using the RNA 6000 Nano assay. Ribosomal RNA was depleted using the NEBNext rRNA Depletion Kit (Bacteria), and RNA-seq libraries made using the NEBNext Ultra™ II Directional RNA Library Prep Kit for Illumina according to manufacturer’s instructions. Library size and adaptor contamination was assessed using the Agilent 2100 Bioanalyser High Sensitivity DNA assay, and concentrations measured with the Qubit dsDNA High Sensitivity assay. Libraries were pooled and for each sample, a minimum of 15 million Paired End 60bp reads were generated on an Illumina NextSeq2000 with unique dual 8 bp indexing. De-duplicated reads were trimmed and their quality assessed using TrimGalore with the --fastqc option (https://github.com/FelixKrueger/TrimGalore). In order to clean the dataset from remnant rRNA reads, trimmed reads were then aligned to the rRNA operons of the NZ_CP064677.1 genome using Bowtie2 (Langmead and Salzberg 2012) and only unmapped reads kept for further analysis. These were mapped onto the NZ_CP064677.1 genome with Bowtie2, and the resulting sam files converted to bam files, sorted and indexed using SAMtools (Li et al. 2009). Read mappings to the *E. coli* and *B. subtilis* genomes, respectively, were counted using the the SAMtools view command and the number of reads mapping to particular genome features (e.g. genes) were computed using the count command of HTseq (Anders et al. 2015). Differential analysis of gene counts was performed using DEseq2 (Love et al. 2014).

Sequencing data for RNA-Seq and Hi-C experiments are available from NCBI GEO under accession GSE341554.

### Comparison to prior transcriptomic data

To compare transcriptional responses in our strains with patterns of differential gene expression observed in prior studies, we made use of a compendium of previously compiled and consistently re-processed RNA-Seq data (Tjaden 2023). We removed datasets not based on total mRNA (e.g. tRNA sequencing) and a large dataset of largely redundant conditions documenting responses to induction of heterologous genes (Garcia et al. 2022). Our own data were re-processed using the same pipeline to render them comparable to the other datasets.

### Growth curves

Precultures of the relevant strains were grown overnight in LB. The OD_600_ of the pre-cultures were adjusted to the lowest one, before being serially diluted to 1:100 in a NUNC96 microplate (ThermoFisher) with LB and a given drug. The microplates were incubated at a given temperature in a Clariostar plate reader (BMG Labtech) for 24h or 48h, with OD_600_ measurement taken every 5 min and orbital shaking at 500 rpm in between measurements.

Doubling time was computed as ln(2)/k where k is the growth rate, defined as (ln(OD_2_) – ln(OD_1_))/(T_2_-T_1_). The time to max slope (i.e. the time before the strain reaches its maximum doubling time) was used a proxy for the duration of the lag phase.

### STED microscopy imaging

Precision microscopy coverslips (Marienfeld) were coated for 15 with a 0.01% dilution of Poly-L-lysine (PLL, Sigma-Aldrich), then rinsed three times with ultrapure water before drying. Exponential phase LB cultures were sampled at an OD_600_ of 0.4 - 0.6, stationary phase LB cultures were sampled from saturated overnight cultures. All cultures were diluted in LB to a total of 2.5 OD_600_ units in 5 ml total. Paraformaldehyde was added to a final concentration of 2%. Samples were fixed for 15 min at 37°C, 180 rpm. The fixation reaction was stopped by addition of glycine to a final concentration of 25 mM. Cells were pelleted for 10 min at 4000 x g and washed in 1X PBS, before being resuspended in 1 ml of 1X PBS. To stain the membrane, NileRed (ChemCruz) was added to a final concentration of 1 µg/ml, to 100 µl of fixed cells. Cells were incubated for 15 min at room temperature in the dark before being washed in 1X PBS. The DNA was stained overnight using Sir-DNA (Spirochrome) at a final concentration of 50 µM. Cells were washed in 1X PBS. The NileRed and Sir-DNA stained cells were resuspended in 300 µl of PBS. 100 µl of the stained cells were pipetted onto PLL-coated coverslips and incubated for 5 min in the dark to let cells adhere. The coverslips were then washed 3 times with 1X PBS before being dried. Coverslips were mounted with Prolong Diamond (Invitrogen) on microscopy slides and left to cure at room temperature in the dark for at least 72h before imaging.

Fluorescence imaging was performed using an inverted TCS SP8 STED 3X (Leica Microsystems, Mannheim, Germany) equipped with a HC PL APO 100x/1,40 OIL STED WHITE immersion objective and running LAS X (version 3.5.10.29396). NileRed and SiR-DNA were excited by 561nm and 633nm laser lines respectively, utilising an 80 MHz pulsed White Light Laser. Fluorescent emission was collected in line sequential mode using Hybrid Detectors (Leica Microsystems, Mannheim Germany) with a spectral window of 570-635nm configured for NileRed, and 650-750nm for SiR-DNA. For confocal imaging, the pixel size was set to 65nm. For 2D STED imaging, a 775nm pulsed laser was used for depletion, and time gating of 0.3-6.0ns applied to both NileRed and SiR-DNA channels, with a pixel size of 20nm and the pinhole closed to 0.5 Airy Units. For 3D STED imaging, the z step size was set to 70nm.

Confocal and STED images were further deconvolved using Huygens Professional software (Scientific Volume Imaging, version 19.10). For confocal and 2D STED images, the Classic Maximum Likelihood Estimation (CMLE) algorithm was utilised, with 40 iterations, and default settings for all other parameters. For 3D STED images, the Good’s roughness Maximum Likelihood Estimation (GMLE) algorithm was used with100 iterations and default settings for all other parameters. MicrobeJ (v5.13o) was used in conjunction with the cell masks to quantify cell morphology.

Cell masks were generated from confocal images using the NileRed fluorescence channel with Omnipose (v1.1.4) (Cutler et al. 2022) and the ‘bact_fluor_omni’ model, applying a flow threshold of 0.5 while retaining default settings for all other parameters.

### Plasmid DNA topology analysis

LB cultures containing 100 µg/ml of ampicillin were inoculated with colonies isolated from a fresh pBR322 transformation. Cultures were processed after reaching 0.4-0.5 of OD_600_. The Monarch Spin Plasmid Miniprep kit (NEB) was used to extract the plasmid following the manufacturer’s protocol with the following modifications. The lysis step was reduced to a maximum of 1 min, all buffers except the elution buffer were used cold and special care was taken to avoid saturating columns. When plasmid yields were too low, the GenElute Plasmid Midiprep kit (Sigma) was used following manufacturer’s instructions. Electrophoresis for 1D and 2D gels was performed as previously described (Villain et al. 2021) with minor modifications. Briefly, 0.8% (for 2D gels) or 0.9% (for 1D gels) agarose was slowly dissolved in 160 ml of 1X TBE buffer. The gels were casted in 14.5 x 14.5 cm trays. At least 1 µg of plasmid was used per condition. Chloroquine was added when required in the first and second dimension to relax the apparent superhelical density of the plasmid and allow proper resolution of the topoisomers. Depending on the agarose and chloroquine concentration, the gels were run at 1.2 V/cm in the first dimension for between 15 and 27 h. For the 2D gels, after being rotated by 90°, the gels were ru at 2 V/cm for 8h. All migrations were performed at room temperature (22°C). Then, gels were washed for at least 15 min in 1X TBE to remove the chloroquine before being stained with SybrGold (Invitrogen) and imaged with a Gel Doc XR+ transluminator (Biorad). Differences in superhelical density between samples were calculated using the equation σ = ΔLk / LK_0_ where LK is the linking number difference of pBR322 between conditions, and LK_0_ the linking number of a relaxed molecule of pBR322. LK was determined using the band counting method (Villain et al. 2021; Keller 1975).

### Nucleoid enrichment and proteomics

Biological duplicates were processed for each condition. Cell pellets from the same culture were used for both the proteomics and RNA-seq. Nucleoid proteins were enriched from gently lysed cells as previously described (Ohniwa et al. 2011; Hocher et al. 2023) with minor modifications. Briefly, cell pellets from exponential or stationary phase cultures were gently resuspended on ice in a mix of 500 µl of buffer A (10 mM Tris-HCl pH 8, 5 mM EDTA, 100 mM NaCl, 10% sucrose), 100 µl of buffer B (100 mM Tris-HCl pH 8.2, 50 mM EDTA, 0.6 mg.ml^−1^ lysozyme) and 10 µl of 100X Halt Protease inhibitor (Thermofisher).

The cells were then lysed for 30 min by addition of 500 µl of buffer C (10 mM Tris-HCl pH 8.2, 10 mM EDTA, 10 mM spermidine, 1% Brij-58 and 0.4% deoxycholate). The lysates were carefully pipetted on 10-50% sucrose gradients, poured per 2 ml layers in ultra-clear 14 × 89 mm tubes (BeckmanCoulter). The samples were centrifuged for 30 min at 12,700 x g in a pre-chilled centrifuge with a SW41-Ti swinging rotor (BeckmanCoulter). The top fraction, enriched for cytosolic proteins (top of the sucrose gradient), and the nucleoid fraction (identifiable as a white viscous band in the middle of the sucrose gradient) were independently collected and processed for methanol/chloroform protein extraction (Wessel and Flügge 1984) . The protein samples were then processed with the iST 8x kit (PreOmics) following manufacturer recommendations.

### Liquid chromatography mass spectrometry (LC-MS) analysis

For each sample 2µg of digest was analysed. Chromatographic separation was performed using an Ultimate 3000 RSLC nano liquid chromatography system (Thermo Scientific) coupled to a Q Exactive HF-X mass spectrometer (Thermo Scientific) via an EASY-Spray source. Electro-spray nebulisation achieved by interfacing to Bruker PepSep emitters (PN: PSFSELJ20, 20µm). Peptide solutions were injected directly onto the analytical column (self-packed column, CSH C18 1.7µm beads, 300μm × 30cm) at a working flow rate of 5 μL/min for 4 minutes. Peptides were then separated using a 70-minute stepped gradient: 0-45% of buffer B for 70 minutes (composition of buffer A – 95/5%: H2O/DMSO + 0.1% FA, buffer B – 75/20/5% MeCN/H2O/DMSO + 0.1% FA), followed by column conditioning and equilibration. Eluted peptides were analysed by the mass spectrometer in positive polarity using a data-independent acquisition mode as follows: an initial MS1 scan was carried out at 120,000 resolution with an AGC target of 3e6 for a maximum IT of 200ms, m/z range: 409.5-1650.5. This was followed by thirty 30K resolution MS2 scans covering 409.5-1650.5 with variable windows sizes set by encyclopeDIA. AGC target set to 3e6 with maximum IT on auto. Normalised collision energy was set to 27%. Total run acquisition time was 80 minutes.

### Proteomics data processing

Data were processed using the Spectronaut software platform (Biognosys, v 19.6.250121.62635) (Bruderer et al. 2015). Pulsar search was performed with default settings for a tryspin/p specific digest with missed cleavage rate set to 3 and a fixed modification of cysteine carbamidomethylation. Variable modifications allowed for methionine oxidation, protein N-terminal acetylation, and cyclisation of glutamine to pyro-glutamate. Searches were carried against a multispecies protein sequence database (downloaded 20240327, 4,191 entries) and a universal protein contaminants database (Frankenfield et al. 2022) (downloaded 08/01/2025, 381 entries). A mutated decoy database approach was employed with PSM, Peptide and Protein group identification FDR = 0.01. Quantification was performed at MS2 level with no value imputation strategy employed and proteotypicity set to “Only proteotypic”. Data have been deposited in the PRIDE database under accession PXD080978.

### Hi-C

Overnight cultures of the relevant *E. coli* strains were diluted to 1/100 and grown in 30 ml LB medium until an OD_600_ of ∼0.4 was reached. Cells were crosslinked with 5% (v/v) formaldehyde (Sigma-Aldrich, cat. no. F8775) for 30 min at room temperature (RT) with gentle agitation (100 rpm). The reaction was quenched by adding 2.5 M glycine (final concentration 0.5 M) for 20 min at RT with gentle agitation, as described in (Cockram et al. 2021). The cells were centrifuged 10 min at 4,000 x g, before to be washed by resuspension in 10 ml of 1X PBS. The cells were centrifuged again before to be resuspended in 1 ml of 1X PBS. Hi-C experiments were performed with a Hi-C kit (Arima Genomics) with a double HpaII + HinfI restriction digestion following manufacturer instructions. Samples were purified using AMPure XP beads (Beckman A63882), suspended in 120ul H_2_O and sonicated in Covaris microTUBE (Covaris, 520045) until average DNA fragment length was approximately 300 bp. Biotinylated DNA was loaded on Dynabeads™ Streptavidin C1 (FISHER SCIENTIFIC, 10202333). Preparation of the samples for paired-end sequencing on an Illumina NextSeq500 (2×35 bp) was performed using Invitrogen TM Collibri TM PS DNA Library Prep Kit for Illumina following manufacturer instructions. All experiments were carried out in biological triplicate.

## CODE & DATA AVAILABILITY

Code and supporting datasets can be found at https://github.com/PaulVillain/NoNAPs_project.

## ACKNOWLEDGEMENTS

We thank Achillefs Kapanidis for discussions and providing access to a Clariostar plate reader and Agnès Thierry for generating Hi-C maps. We further thank Peter Sarkies, Dave Sherratt, and Rob Klose for comments on an earlier version of the manuscript. This work was made possible by funding from the UKRI Medical Research Council (MC-A658-5TY40), the John Fell Fund (ALD00350) and the Department of Biochemistry, University of Oxford (to TW) and a grant from the French government, managed by the Agence Nationale de la Recherche under the France 2030 program (ANR-23-CHBS-0002) to RK. PV is supported by a Wellcome Trust Early Career Award (324805/Z/25/Z).

## SUPPLEMENTARY TABLES

**Table S1.** Strains used and generated in this study

**Table S2.** Primers used in this study

**Table S3.** Mutations in ΔNAP9

**Table S4.** Spearman correlation coefficients and corresponding P values between transcriptional profiles of WT, ΔNAP5, ΔNAP9 and public data. P values <1e-200 are given as 0.

## CONFLICT OF INTEREST

The authors declare that no conflict of interest exists.

## Notes

### Competing Interest Statement

The authors have declared no competing interest.

