## Supplementary material for "Chromatin is dispensable for bacterial life": Table S1

**Table S1. Strains used and generated in this study**

| **Strain name** | **Genotype** | **Markers** | **Source** |
| --- | --- | --- | --- |
| *E. coli* XL1 Blue | *endA*1, *gyrA*96, thi-1, *recA*1, *relA*1, lac *glnV*44, F'[ ::Tn10, *proAB+, lacIq,* Δ(l*acZ*)M15], *hsdR*17 | Tetracycline resistance Nalidixic acid resistance | Stratagene |
| *E. coli* BW25113 (WT) | F^-^, LAM^-^, rrnB3, DElacZ4787, hsdR514, DE(araBAD)567, DE(rhaBAD)568, rph-1 | / | Lab collection |
| *B. subtilis* 168 | *trp*C2 | Tryptophane auxotrophy | Lab collection |
| *E. coli* JW3964 (Keio) | BW25113, ∆*hupA* | Kanamycin resistance | Lab collection |
| *E. coli* JW0430 (Keio) | BW25113, ∆*hupB* | Kanamycin resistance | Lab collection |
| *E. coli* JW1225 (Keio) | BW25113, ∆*hns* | Kanamycin resistance | Lab collection |
| *E. coli* JW1702 (Keio) | BW25113, ∆*ihfA* | Kanamycin resistance | Lab collection |
| *E. coli* JW0895 (Keio) | BW25113, ∆*ihfB* | Kanamycin resistance | Lab collection |
| *E. coli* JW2644 (Keio) | BW25113, ∆*stpA* | Kanamycin resistance | Lab collection |
| *E. coli* JW0797 (Keio) | BW25113, ∆*dps* | Kanamycin resistance | Lab collection |
| *E. coli* JW3229 (Keio) | BW25113, ∆*fis* | Kanamycin resistance | Lab collection |
| *E. coli* JW0872 (Keio) | BW25113, ∆*lrp* | Kanamycin resistance | Lab collection |
| *E. coli* ∆NAP1 | BW25113, ∆*hupA* | / | This study |
| *E. coli* ∆NAP2 | BW25113, ∆*hupA*, ∆*hupB*, *rpsB* (K132) | / | This study |
| *E. coli* ∆NAP3 | BW25113, ∆*hupA*, ∆*hupB*, ∆*hns*, *rpsB* (K132) | / | This study |
| *E. coli* ∆NAP4 | BW25113, ∆*hupA*, ∆*hupB*, ∆*hns*, ∆*ihfB*, *rpsB* (K132), *mnmE* (A310T) | / | This study |
| *E. coli* ∆NAP5 | BW25113, ∆*hupA*, ∆*hupB*, ∆*hns*, ∆*ihfB*, ∆*ihfA*, *rpsB* (K132), *mnmE* (A310T), *topA* (∆RET 790-792), *proV* (P197L) | / | This study |
| *E. coli* ∆NAP6 | BW25113, ∆*hupA*, ∆*hupB*, ∆*hns*, ∆*ihfB*, ∆*ihfA*, ∆*stpA* BW25113, ∆*hupA*, ∆*hupB*, ∆*hns*, ∆*ihfB*, ∆*ihfA*, *rpsB* (K132), *mnmE* (A310T), *topA* (∆RET 790-792) | / | This study |
| *E. coli* ∆NAP7 | BW25113, ∆*hupA*, ∆*hupB*, ∆*hns*, ∆*ihfB*, ∆*ihfA*, ∆*stpA*, ∆*dps* BW25113, ∆*hupA*, ∆*hupB*, ∆*hns*, ∆*ihfB*, ∆*ihfA*, *rpsB* (K132), *mnmE* (A310T), *topA* (∆RET 790-792) | / | This study |
| *E. coli* ∆NAP8 | BW25113, ∆*hupA*, ∆*hupB*, ∆*hns*, ∆*ihfB*, ∆*ihfA*, ∆*stpA*, ∆*dps*, ∆*fis* BW25113, ∆*hupA*, ∆*hupB*, ∆*hns*, ∆*ihfB*, ∆*ihfA*, *rpsB* (K132), *mnmE* (A310T), *topA* (∆RET 790-792), *dnaA* (R302S) | / | This study |
| *E. coli* ∆NAP9 | BW25113, ∆*hupA*, ∆*hupB*, ∆*hns*, ∆*ihfB*, ∆*ihfA*, ∆*stpA*, ∆*dps*, ∆*fis*, ∆*lrp*, *rpsB* (K132), *mnmE* (A310T), *topA* (∆RET 790-792), *dnaA* (R302S),  *ymfE*-*lit* intergene | / | This study |
