## Supplementary material for "Chromatin is dispensable for bacterial life": Table S3

**Table S3. Mutations in** ∆**NAP9**

| **Mutations in** ∆**NAP9** | **Genomic alteration** |
| --- | --- |
| *rpsB* (K132) | 186753: G 🡪 GAAA |
| *mnmE* (A310T) | 3881115: G 🡪 A |
| *topA* (∆RET 790-792) | 1327669: TCGCGTGAAA 🡪 T |
| *proV* (P197L) | 2798763: C 🡪 T |
| *dnaA* (R302S) | 3876186: G 🡪 T |
| *ymfE*-*lit* intergene | 1193909: AATGAAATG 🡪 A |
