## Supplementary material for "Chromatin is dispensable for bacterial life": Figures S1-8

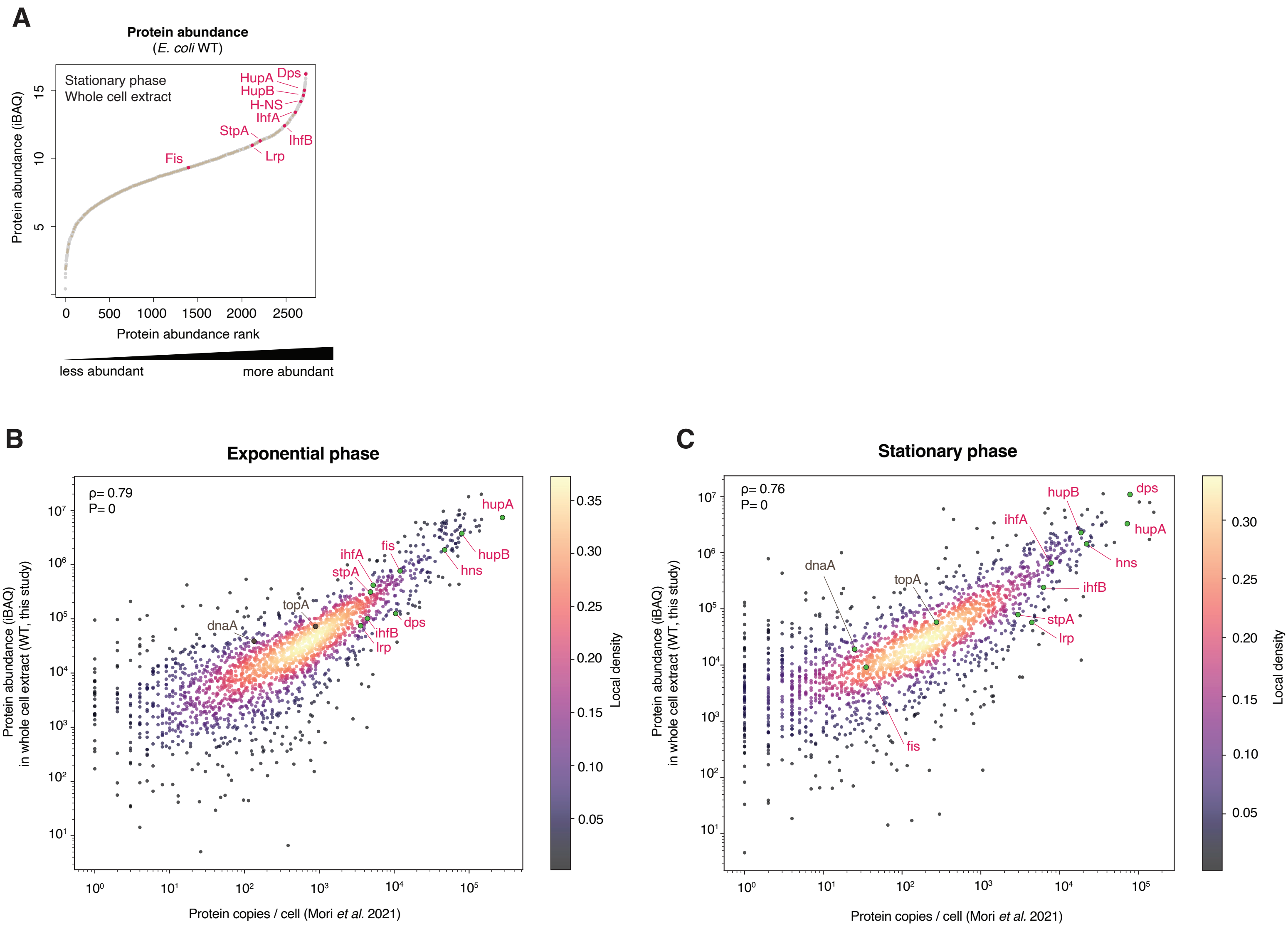


**Figure S1. Protein abundance in ΔNAP9. A.** Ranked protein abundance in WT *E. coli* cells during stationary phase, as determined using quantitative mass spectrometry. NAPs (pink) are highlighted along with select other proteins. This panel corresponds to Figure 1C, which shows ranked abundance in exponential phase. **B,C.** Protein abundance in WT cells in exponential (B) an stationary (C) phase compared to prior data quantifying protein copies per cell. NAPs (pink) are highlighted along with DnaA and TopA.
iBAQ: intensity-based absolute quantification.

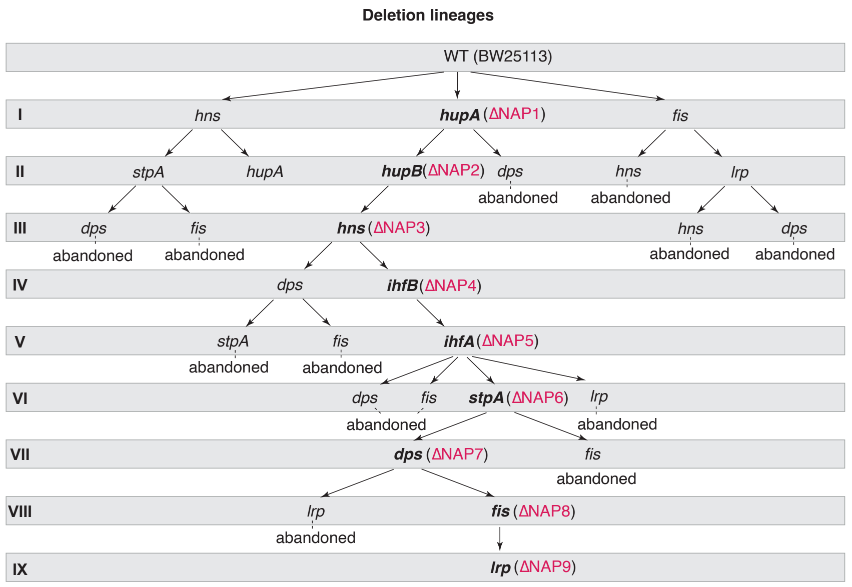


**Figure S2. Order of deletion leading up to ΔNAP9 and deletion series abandoned along the way.**

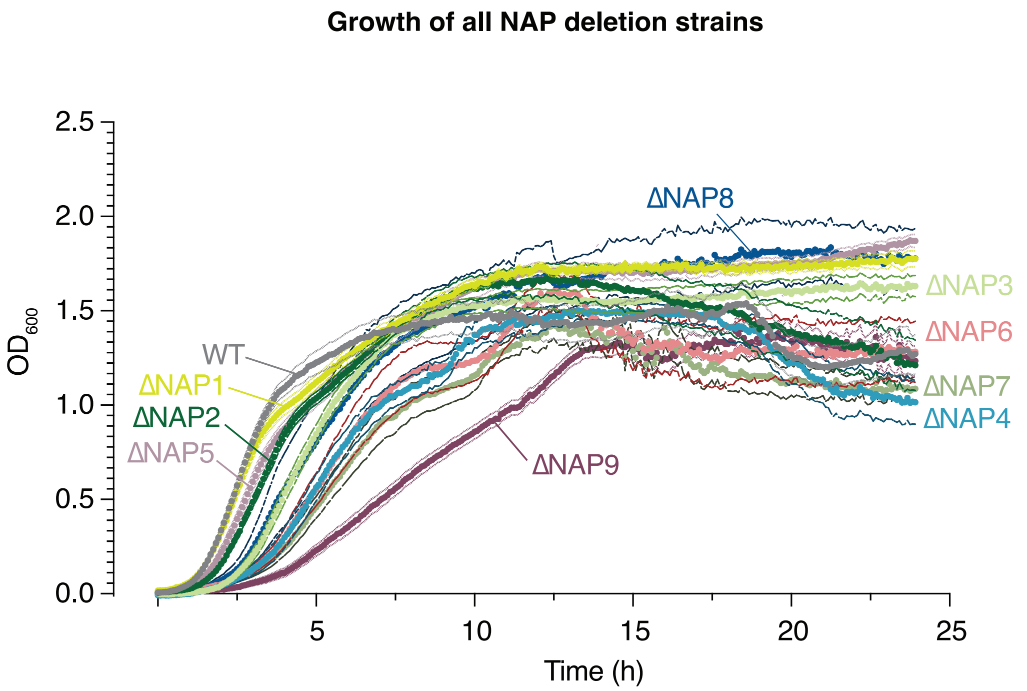


**Figure S3. Growth of intermediate deletion strains.** Growth curves of WT, ΔNAP9, and all intermediate strains (ΔNAP1-ΔNAP8), based on 3 biological replicates. For each strain, the central line indicates the mean across 3 biological replicates, the thinner outer lines represent the SEM.

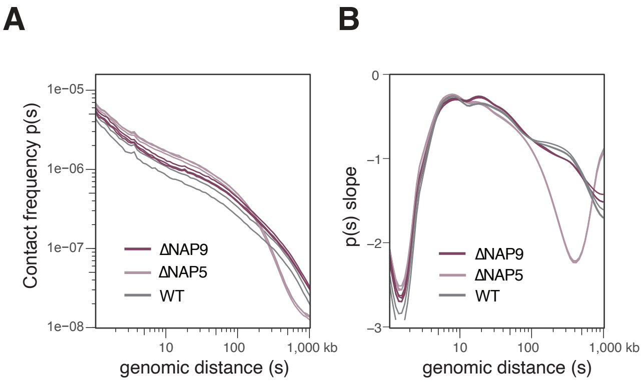


**Figure S4. Additional feature of Hi-C maps. A,B.** Contact frequency probability (A) and slope of the contact frequency probability (contact decay) (B) extracted from Hi-C matrices of WT, ΔNAP5 and ΔNAP9. Three biological replicates for each condition are shown.

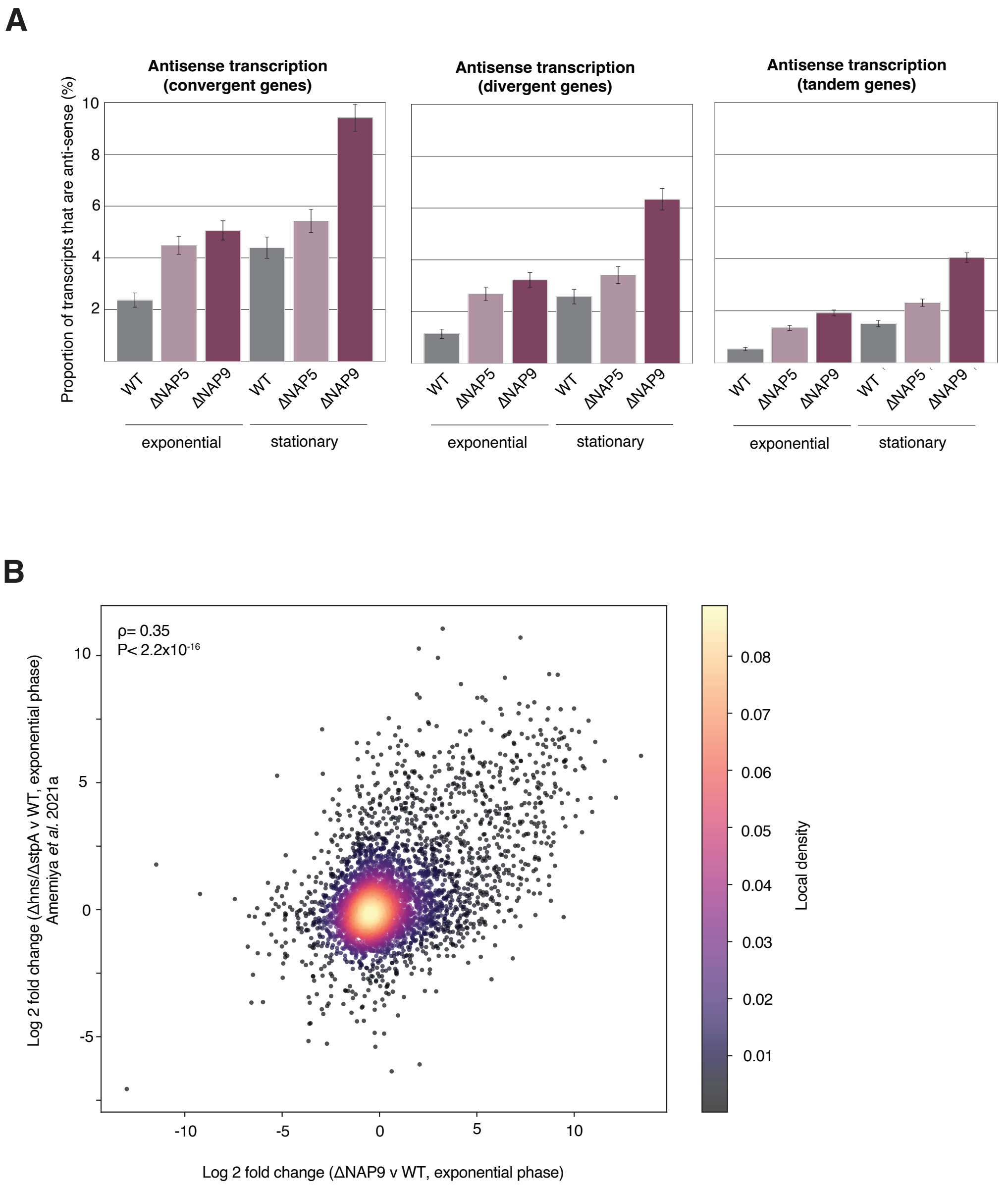


**Figure S5. Transcriptional dysregulation in ΔNAP9. A.** Proportion of sequencing reads mapping antisense to known transcription units in WT, ΔNAP5 and ΔNAP9, split according to whether the focal gene and its neighbour are convergently transcribed (left panel), divergently transcribed (middle panel) or oriented in tandem (right panel). This analysis demonstrates that increased antisense transcription is not explained by higher reads of readthrough from divergently oriented neighbouring gene. Error bars indicate SEM. **B.** Correlation between the changes observed in ΔNAP9 (vs WT) compared to Δhns/ΔstpA double deletion (vs WT), highlighting shared response amongst strongly upregulated genes normally repressed by H-NS/StpA.

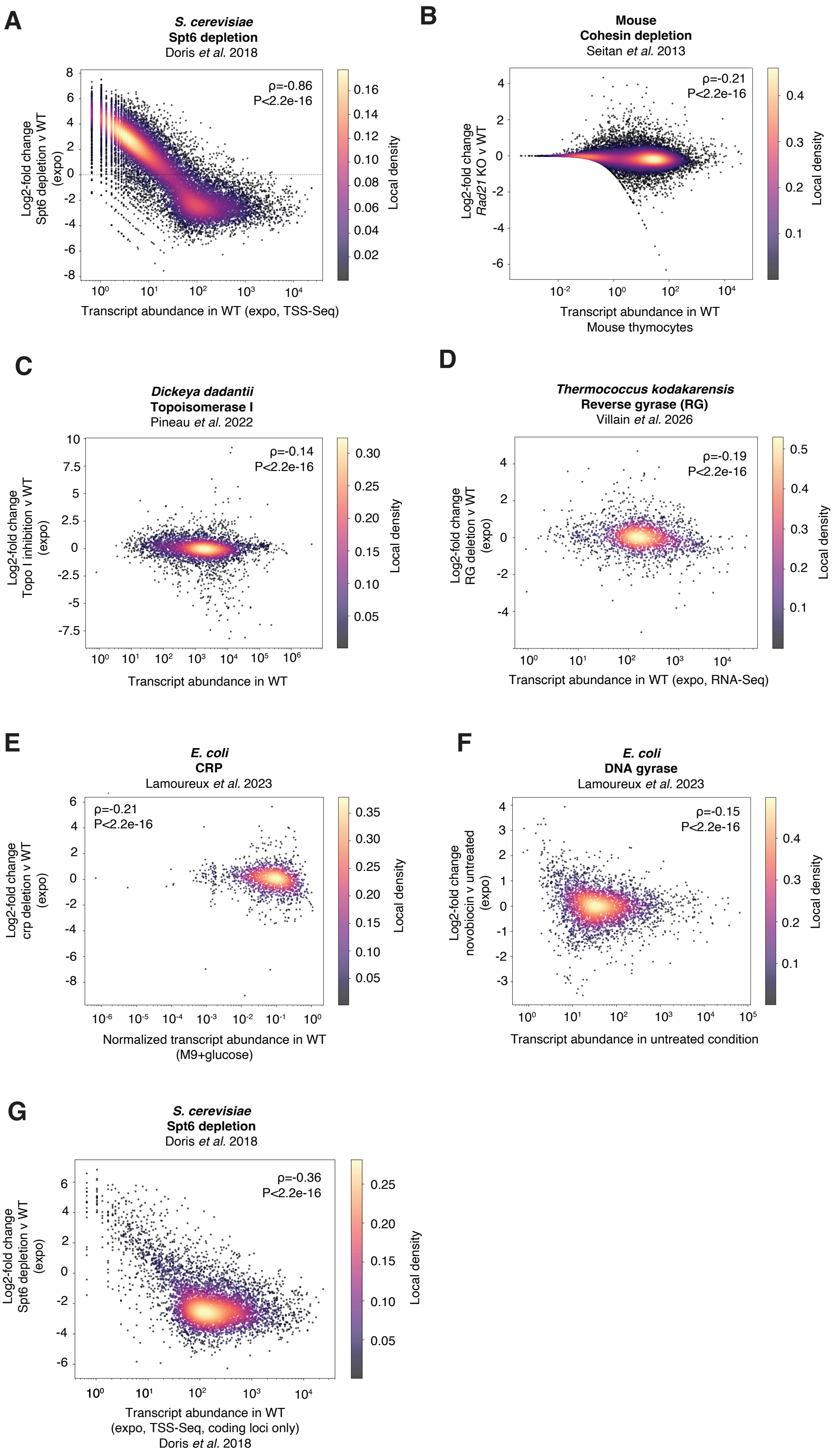


**Figure S6. Global homogenization of gene expression following interventions involving global regulators of transcription.**
**A-G.** Transcriptional changes following a given intervention as a function of transcript abundance in the corresponding wildtype/control. **A.** Spt6 depletion in *S. cerevisiae*, this is the same panel as Figure 3J and is given for comparison **B.** cohesin depletion in mouse thymocytes, **C.** inhibition of topoisomerase I in *D. dadantii*, **D.** deletion of reverse gyrase (RG) in *T. kodakarensis*, **E.** deletion of *crp* in *E. coli*, **F.** inhibition of DNA gyrase in *E. coli*, **G.** Spt6 depletion in *S. cerevisiae*, showing TSS-seq data mapping to the promoters of coding genes only.

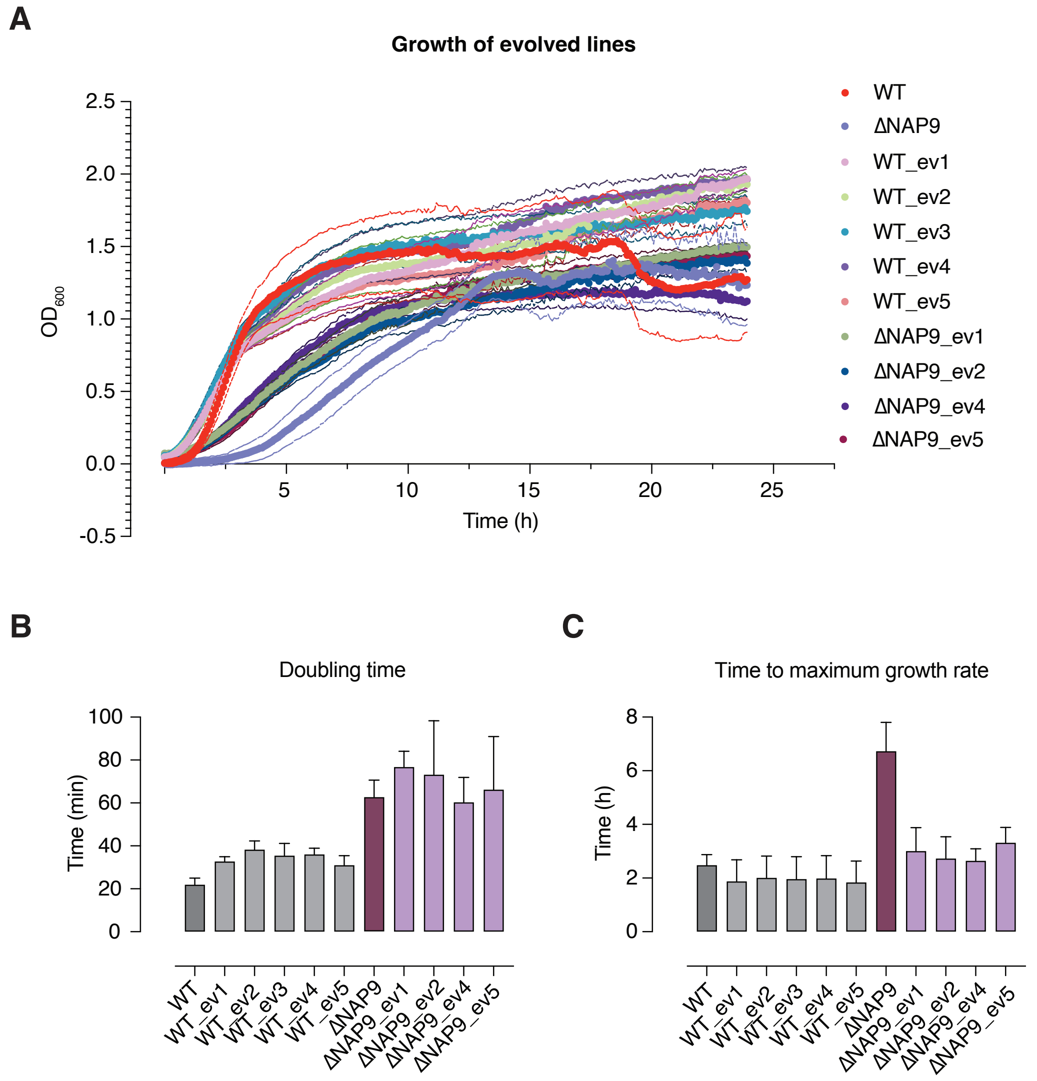


**Figure S7. Growth of evolved lines A.** Growth curves of evolved lines based on 3 biological replicates. For each strain, the central line indicates the mean across 3 biological replicates, the thinner outer lines represent the SEM. WT and ΔNAP9 ancestors are given as reference. **B, C.** Doubling time (B) and time to maximum growth rate (C) based on the growth curves in A.

A

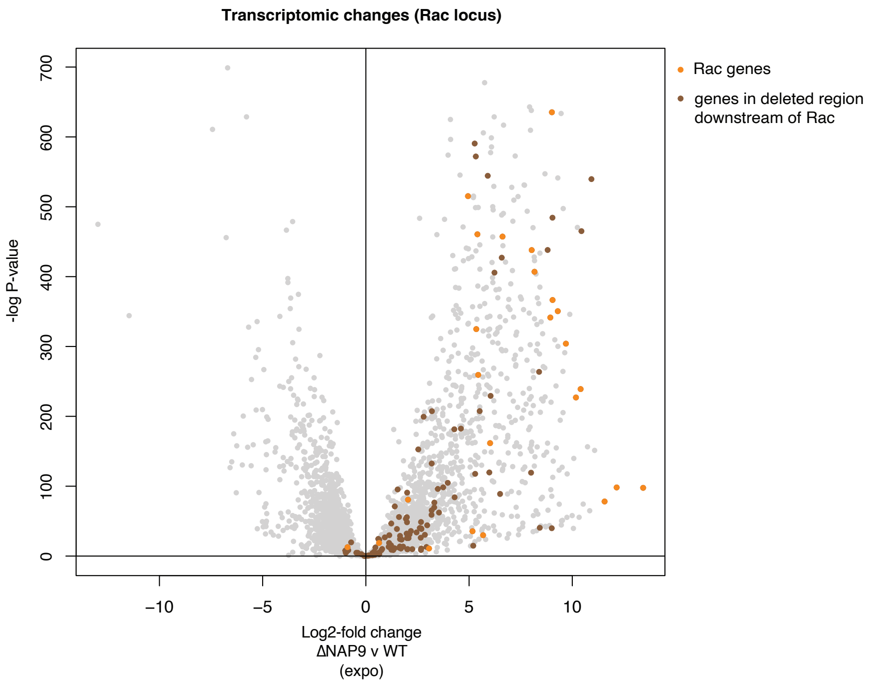


B

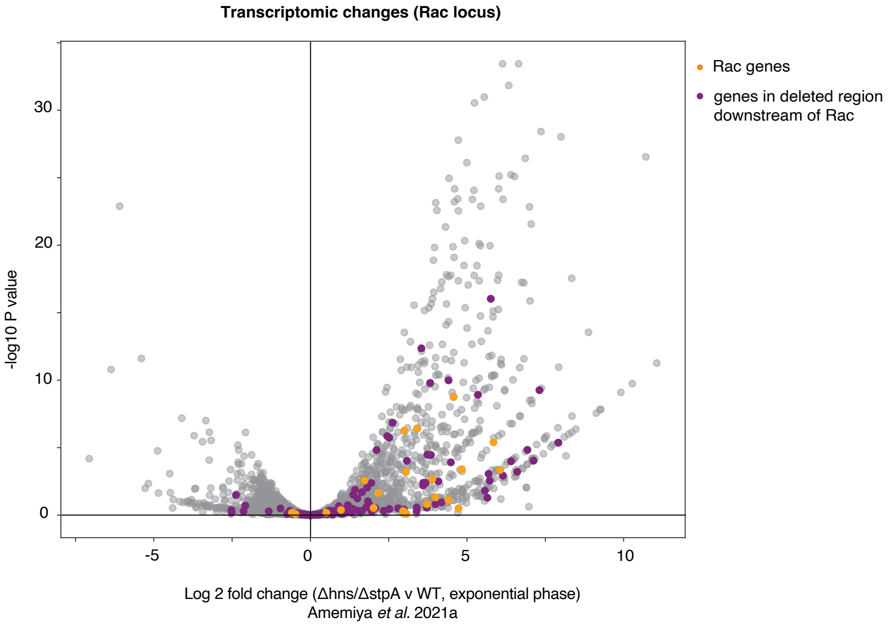


**Figure S8. Transcriptional changes around the Rac locus A,B.** Changes in RNA abundance in ΔNAP9 (vs WT) (A) and the Δ*hns*/Δ*stpA* double deletion (vs WT) (B), highlighting changes experienced by genes encoded by the cryptic prophage Rac (orange), and genes downstream of Rac (brown/purple), in the region that has been recurrently deleted in lines evolved from ΔNAP9 (see Figure 5).
